# Trace elements in grey seals from the Gulf of St. Lawrence

**DOI:** 10.1101/2021.08.30.458200

**Authors:** Gwyneth A. MacMillan, Marc Amyot, Pierre-Yves Daoust, Mélanie Lemire

## Abstract

We measured baseline levels of 19 trace element and mercury speciation for grey seals (*Halichoerus grypus*) from the Gulf of St. Lawrence (GSL), Québec, Canada. With interest growing in commercializing grey seal products for human consumption in this region, the goal of this study was to measure essential and non-essential trace elements in grey seals to evaluate health concerns and nutritional benefits. From 2015 to 2019, 120 grey seals were sampled by hunters and researchers at 4 sites in the GSL. Muscle, liver, heart and kidney samples were analyzed for 10 non-essential elements (Sb, As, Be, B, Cd, Pb, Hg, Ni, Tl, Sn) and 9 essential elements (Co, Cr, Cu, Fe, Mg, Mn, Mo, Se, Zn). Both total mercury (THg) and methylmercury (MeHg) were analysed for a subset of samples. Many elements were undetected in liver (Sb, As, Be, B, Cr, Co, Pb, Ni, Tl, Sn) and muscle tissues (same, plus Cd, Mn, Mo). Results showed lower element concentrations in the muscle (Fe, Mg, Se) and livers (Cd, Cr, Hg, Mn, Mo, Se) of young-of-the-year harvested in the winter (< 6 weeks old) compared to older animals feeding at sea. For older seals (∼ 5 months to 29 years), we did not observe progressive age-dependent bioaccumulation. Sex-specific differences were not very pronounced, but a few elements were 30 - 70% higher in the muscle (THg, MeHg) and liver (Mn, Zn) of male seals. Comparison to Canadian dietary reference intakes shows that a weekly portion of liver from young-of-the-year (< 6 weeks old) is a good source of essential elements (Cu, Fe) and that muscle and liver from this age category does not exceed reference values for toxic elements (As, Cd, Pb, MeHg). Ongoing discussions with regional public health professionals will help to develop dietary recommendations for the consumption of older grey seals.

**HIGHLIGHTS:**

1. We measured baseline levels of 19 trace elements in grey seals harvested from the Gulf of St. Lawrence.
2. We evaluated nutritional benefits and health concerns of human consumption of grey seal products.
3. Once seals began feeding at sea (∼ 5 mo), many element concentrations increased, but did not bioaccumulate progressively with age afterwards.
4. Some elements were more concentrated in the muscle (mercury) and livers (manganese, zinc) of male seals.
5. Young seal (< 6 we) livers are a good dietary sources of copper and iron, while its muscle and liver were below reference values for toxic elements.

**GRAPHICAL ABSTRACT:** 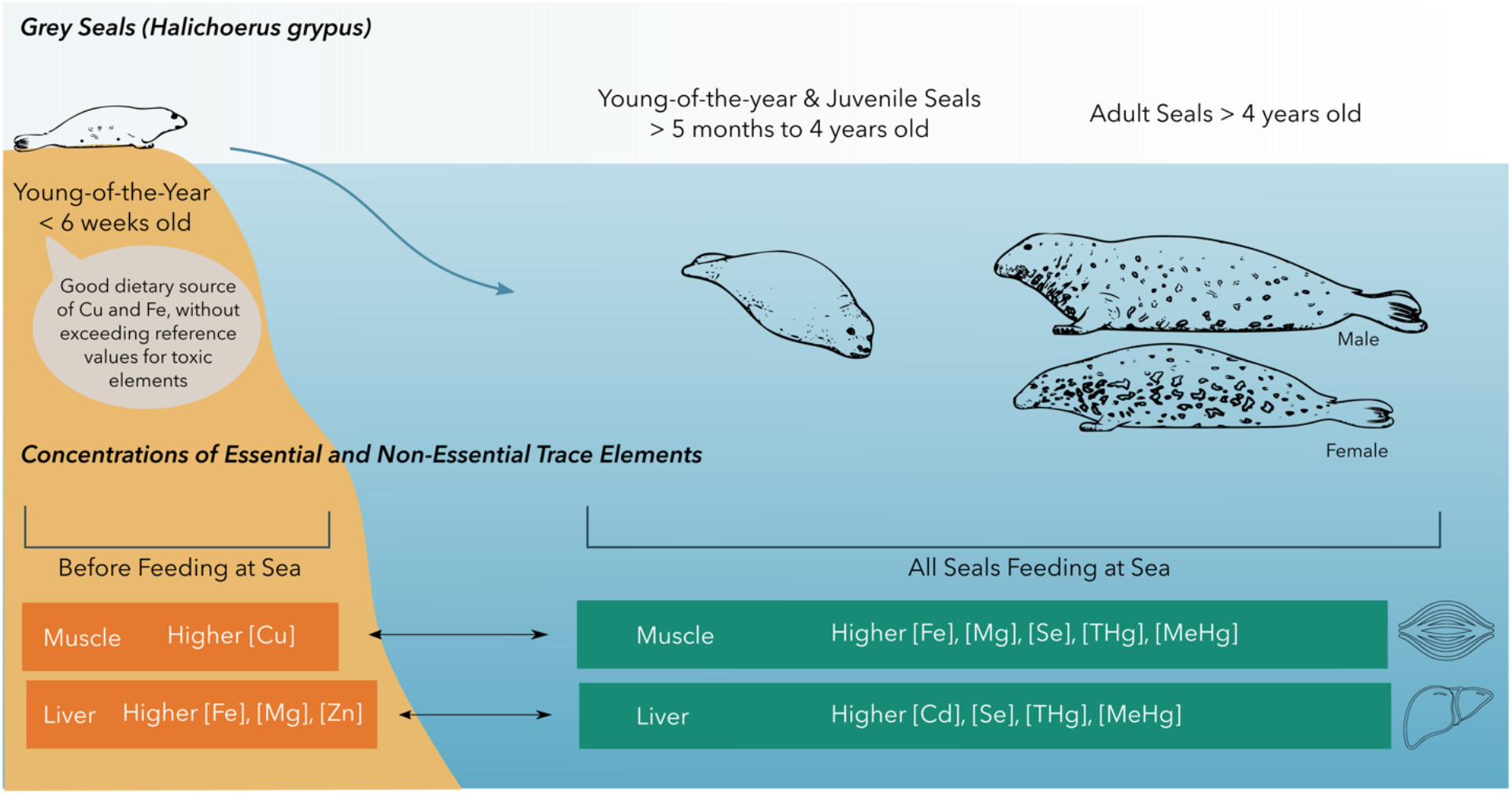

## 1. INTRODUCTION

There is currently growing interest in the commercial exploitation of grey seals (*Halichoerus grypus)* from the Gulf of St. Lawrence, Canada. Seal hunting is a cultural tradition in the region and an important economy for many fishermen in the Magdalen Islands, Québec, located in the Gulf of St. Lawrence (GSL) (Pintal, 2003; Baril and Breton, 1982). Until the 19th century, Magdalen Islanders practiced a subsistence seal hunt for meat, oil and clothing, and, today, small businesses have developed new uses and markets for grey seal products with the goals of promoting local benefits and sound resource management (ACPIQ, 2021). Grey seals in eastern Canadian waters compose a single genetic population estimated to number 424,300 individuals in 2016 and distributed across 3 breeding areas: Sable Island off Nova Scotia, coastal areas of Nova Scotia, and the GSL (Hammill et al., 2017). Since the end of an intensive hunt in the 1960s, the population has been slowly increasing and increased predation by grey seals may be impacting the recovery of cod and other groundfish stocks which collapsed due to overfishing (Hammill et al., 2014; Swain et al., 2019)). The total allowable catch for grey seals in Canada is set at 60,000, however only 1,612 seals were harvested commercially in 2016 which is less than 1% of the Northwest Atlantic population (Government of Canada, 2016a). The Committee on the Status of Endangered Wildlife in Canada (COSEWIC) has designated the grey seal as “not at risk” since 1999.

In the Northwest Atlantic, grey seal pups are born between December and February on land or ice. Unlike harp seals (*Pagophylus groenlandicus*), grey seals do not need access to pack ice to give birth. Most pups (85%) are born on Sable Island, and the rest are born in the GSL (11%) and along the Eastern Shore of Nova Scotia (4%). The distribution of pups has changed over time, with a decline in the proportion born on ice in the GSL and an increase on Sable Island and the Eastern Shore (Government of Canada, 2016b). Pups are weaned at about 2-3 weeks but remain after on land until about 40 days, when they head out to sea to feed after a post-weaning fast (Noren et al., 2008). Once at sea, grey seals are mobile, generalist predators that feed on demersal fish such as cod, American plaice, and sand lance, as well as small pelagic fish such as herring and mackerel. Grey seal diets can vary by geography, season, and sex (Hammill et al., 2014). Grey seals seem to occupy an intermediate trophic level compared to other seals and marine mammals in the Estuary and GSL. Previous studies indicate that hooded seals (*Cystophora cristata*) occupy a higher trophic level than grey seals, whereas harp seals and beluga whales (*Delphinapterus leucas*) in the region occupy lower trophic levels (Lesage et al., 2001; Morissette et al., 2006).

Commercial hunts for grey seals are currently centered around the Magdalen Islands and in the southern GSL and primarily target post-weaned young-of-the-year (YY) less than 6 weeks old for meat, fat, and internal organs, although juvenile and adult seals are also sometimes hunted for meat. For predators at the top of marine food webs, such as seals, the processes of bioaccumulation and biomagnification can lead to higher concentrations of some trace elements (AMAP, 2018). Essential trace elements, such as copper and iron, are necessary for the proper biological functioning of organisms but can reach high concentrations harmful to the health of organisms or those who consume them. Non-essential trace elements, such as mercury, cadmium and lead, have no biological function and can be toxic at very low concentrations for human consumers, especially during pregnancy and for young children (Fraga, 2005).

To evaluate possible nutritional benefits and health concerns related to the human consumption of grey seal products, the objective of this study was to measure baseline concentrations of several essential and non-essential trace elements in grey seals harvested from the GSL. For this study, 120 grey seals of different ages were sampled from 2015 to 2019 at four different sites in the GSL. More specifically, we wanted to evaluate the i) influence of age category (YY, juvenile, adult), exact age (years), sex, sampling site and tissue type (muscle, liver, kidney, heart) on trace element concentrations and ii) compare values measured in grey seals with Canadian dietary reference intakes and reference values for essential and non-essential elements respectively.

## 2. MATERIAL & METHODS

### 2.1 Sample collection

From 2015 to 2019, 120 grey seals were harvested during commercial or scientific sampling at two sites near the Magdalen Islands (47.3877° N, 61.9012° W) and two sites in the Northumberland Strait (45.9713° N, 63.5608° W) in the GSL, Canada (Fig. 1). Grey seals harvested during scientific sampling were collected under permits QUE-Research Notice IML 2015-19, IML 2017-01, and QUE-SCIENTIFIQUE-069-2018. The sampling during commercial hunts were collected under Department of Fisheries and Oceans (DFO) permits IM 2015-01, IM 2017-01, and IM 2018-01. Sample handling was authorized by the UPEI Biosafety Committee (protocol 6006710). Sample collection was coordinated by the *Association des chasseurs de phoques Intra-Québec (ACPIQ)*, the *Ministère de l’Agriculture, des Pêcheries et de l’Alimentation du Québec* (MAPAQ) and the Maurice-Lamontagne Institute of Fisheries and Oceans Canada.

**Figure 1.**
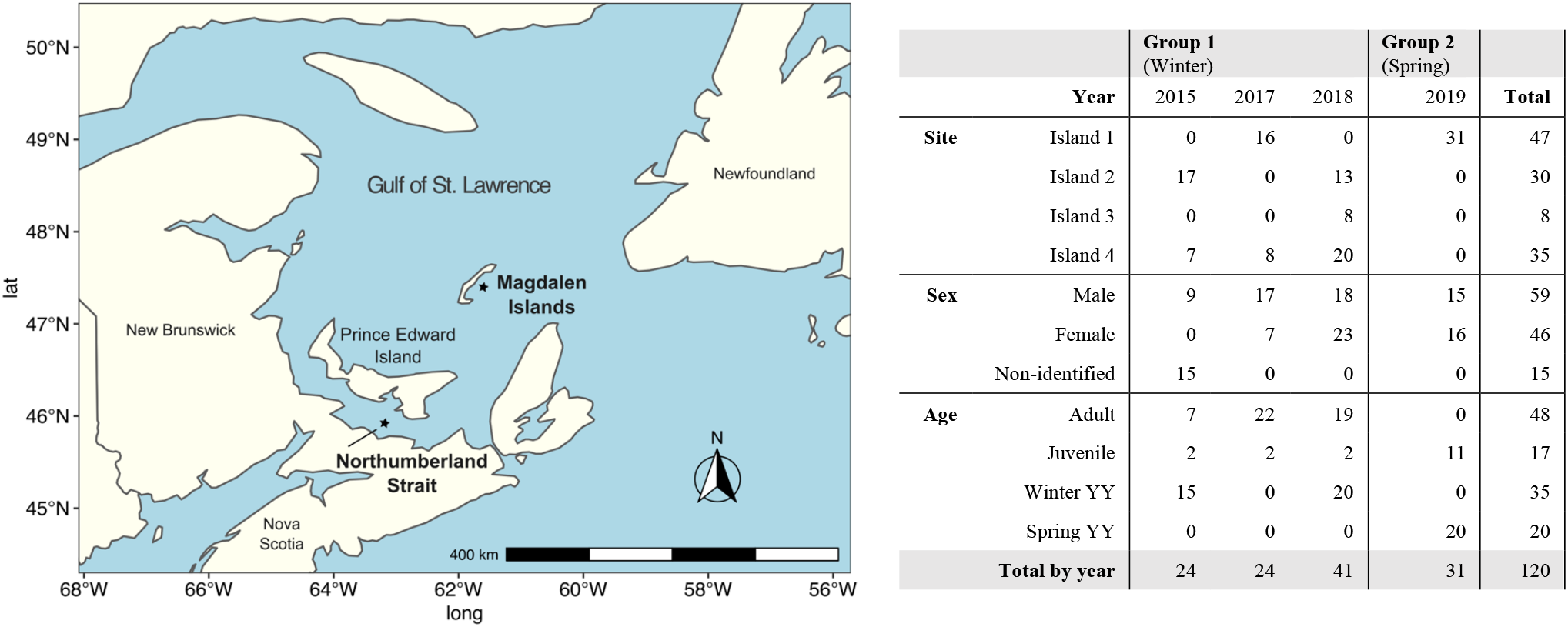
Left: Map showing grey seal sampling sites in the Gulf of the St. Lawrence (n = 120). Islands 1 and 2 are located near the Magdalen Islands and Islands 3 and 4 are location in the Northumberland Strait. Right: Table of the number of grey seals sampled by year, site, sex and age category. Trace elements were analyzed in muscle and liver tissues for Group 1 and in muscle, heart and kidney tissues for Group 2.

Samples were collected in two distinct groups. The first group (Group 1) consisted of 89 seals harvested in January and February of 2015, 2017 and 2018 at four sites designated Islands 1 to 4. Note that Island 4 was sampled as a comparison and no commercial seal hunting is conducted at this location. Only muscle and liver tissues were collected from this group of seals which included 36 winter YY, 6 juveniles, and 48 adults (Fig. 1). Muscle tissue was collected from inside the abdomen to avoid contamination from bullet fragments, except for samples from 2015 which were taken from several places on the carcass. A second group (Group 2) of 31 samples were collected from sub-adult seals harvested near Island 1 in June 2019. Muscle, heart and kidney (but not liver) tissue samples were collected from this group of 20 spring YY and 11 juveniles. For both groups, about 50 g of each tissue were collected from each seal. All samples were kept cold during transport and were frozen at -20°C less than 24 hours after collection.

During commercial expeditions, recently weaned young-of-the-year (< 6 weeks) were harvested on dry land using a hakapik and older seals (> 5 months) were harvested in the water using a high caliber rifle from a small motorboat. During scientific expeditions, adult seals were also harvested on dry land using a high caliber rifle. Sex was determined at the time of sampling and seals were initially classified into 3 age categories based on size and fur appearance; young- of-the-year (YY, < 1 year), juvenile (1 - 4 years) or adult (> 4 years). The exact age was estimated for a sub-sample of seals (n = 54) by counting the annual growth layers in the cementum of canine teeth (Frie et al., 2013). Due to the difference in sampling dates (winter vs. spring) for the two groups, seals were subsequently grouped into 4 age categories based on sampling date and canine analysis; winter young-of-the-year (winter YY, < 6 weeks old), spring young-of-the-year (spring YY, 5-6 months old), juvenile (1 - 4 years), or adult (> 4 years).

### 2.2 Laboratory analysis

For Group 1, muscle (n = 88) and liver (n = 89) samples were analyzed for 9 essential trace elements: chromium (Cr), cobalt (Co), copper (Cu), iron (Fe), magnesium (Mg), manganese (Mn), molybdenum (Mo), selenium (Se), zinc (Zn), and 10 non-essential elements: antimony (Sb), arsenic (As), beryllium (Be), boron (B), cadmium (Cd), lead (Pb), mercury (Hg), nickel (Ni), thallium (Tl), and tin (Sn). For Group 2, muscle, heart and kidney samples (n = 31) were analyzed for the same 19 elements. Trace elements were analyzed at the Animal Health Laboratory, Laboratory Services Division, University of Guelph, Ontario, Canada. Complete samples (about 50 g) were cut in half and a sub-sample of tissue was taken from the freshly exposed portion to avoid contamination from outside surfaces. Non-homogenized wet tissue samples (about 1 g) were weighed and digested with nitric acid in a microwave (CEM MARSXpress) and then diluted with Nanopure water before analysis by Inductively Coupled Plasma - Mass Spectrometry (ICP-MS). Detection limits (LOD) and detection frequencies (percentage of samples > LOD) are shown in Table S1. Note that LOD were lower for samples from Group 2 because the two groups of samples were analysed on different instruments (Group 1 - 820 MS, Varian Inc. Palo Alto, CA, USA; Group 2 - 7900 ICP-MS, Agilent, Santa Clara, CA, USA).

Total mercury (THg) and methylmercury (MeHg) were analysed for a sub-sample from Group 1 (muscle : n = 32, liver : n = 43) and all samples from Group 2 (muscle, heart and kidney : n = 31). Analyses were conducted at the laboratory of the *Chaire de recherche du Canada en écotoxicologie et changements mondiaux*, Université de Montréal, Québec, Canada (Table S1). Tissue samples were freeze-dried and homogenized before analysis following trace metal clean protocols. Tissue THg was analyzed using two different methods. The least concentrated samples (all muscle, heart, and kidney tissues, as well as liver tissue from young-of-the-year) were analyzed by cold vapour atomic absorption spectrometry with a Direct Mercury Analyzer (DMA-80, Milestone Inc, Shelton, CT, USA). The more concentrated samples (all liver samples, except for young-of-the-year) were analyzed using cold-vapor atomic fluorescence spectrometry (CV-AFS). For this analysis, samples were diluted and analyzed with a Tekran 2600 using EPA Method 1631. Tissues MeHg was analyzed by CV-AFS with a Tekran 2700 using EPA method 1630 (Tekran Instruments Corporation, Seattle, WA, USA). Concentrations were converted from dry weight to wet weight using the percent humidity of each sample. Percent humidity was 70 ± 2.6 %, 70 ± 1.9 %, 77 ± 0.9 %, and 77 ± 0.9 % (mean ± SD) for samples of muscle, liver, heart and kidney respectively.

Analytical blanks, duplicate samples, an internal control (bovine liver), and a certified reference material (Tort-3 - lobster hepatopancreas; NRC Canada) were included after each set of 40 samples during trace element analysis at the University of Guelph (Table S2). Internal or certified standards were used to validate instrument drift during analysis. Acceptable limits for duplicate samples were 20% for elements measured at more than twice the LOD and recovery varied from 2 to 16%. For the THg and MeHg analyses at the *Université de Montréal*, analytical blanks, internal controls (CALA-4 - water; Canadian Association for Laboratory Accreditation), and certified reference materials (Tort-2 - lobster hepatopancreas, Dorm-4/Dorm-2 - fish tissue, and SO-2 - soil; NRC Canada) were analysed after each 10-12 samples. Reference material measurement fell within the range of certified values for all analyses (Table S3). A sub-sample of muscle (n = 32) and liver (n = 45) samples from Group 1 and all samples from Group 2 (n = 31) were analyzed for THg at both laboratories in this study. Correspondence between analyses was acceptable with an average difference of 4.9% (Table S4). To maximize sample size in this study, all reported values for THg are taken from analyses by ICP-MS at the University of Guelph. One exception to this was when reporting MeHg values, which were compared to the corresponding THg value from the same analysis batch at the *Université de Montréal* to improve the accuracy of measurements.

### 2.4 Data handling and statistical analysis

Trace element concentrations were not normally distributed and therefore geometric means (GM), which are the antilogarithm of the mean of the logarithmic values for each sample, were used to show the central tendency of the data. The 95% confidence intervals (95% CI) are also reported. All concentrations are shown on a wet weight (w.w.) basis. Sample concentrations under the LOD were replaced with half the LOD. An analyte was considered “detected” if more than 60% of samples were higher than the detection limits (LOD) of the instrument, and GM and 95% CI were only reported for detected elements.

One-way Student’s t-tests (independent samples) and one-way analysis of variance (ANOVA) followed by Tukey HSD post-hoc tests were used to evaluate differences in trace element concentrations independently by tissue type, age category, sex, and sample site. Pearson’s r correlations were used to determine the relationship between variables, including exact age (years) vs. trace element concentrations and Se vs. THg concentrations. Unbalanced two-way ANOVAs were used to test the relative effect of sex and age category on THg and MeHg concentrations in muscle and liver tissues. All statistical tests were performed using log_10_- transformed concentrations and a threshold of α equals 0.05 in “R” software (R Core Team, 2020). Graphics were created with the ‘*ggplot2*’ package (Hadley, 2016).

Concentrations of non-essential trace elements were compared to reference values established by Health Canada for chemical contaminants in food (Canada, 2005a). Reference values are intended to prevent excessive exposure to contaminants for the most vulnerable populations, usually pregnant or nursing women and young children, while maximizing the health benefits of consuming the food item. For mercury, the reference values of 0.5 and 1.0 µg Hg/g for commercially sold fish products are calculated from the provisional Tolerable Daily Intake (pTDI) of 0.47 µg Hg/kg of body weight for the general population and of 0.2 µg Hg/kg of body weight for young children and pregnant women (including women of childbearing age, those who plan to become pregnant, and breastfeeding women). This correspond to dietary recommendations of a maximum of 150 g per week for pregnant women and 75 g per week for children aged 1 to 4 years, equivalent to 1 or 2 portion sizes, of a food item with about 0.5 µg Hg/g (but below the reference value of 1.0 µg Hg/g) (Canada, 2007).

Concentrations of essential trace elements were compared to Health Canada’s Dietary Reference Intake Tables (Canada, 2005b) using four scenarios for two population subgroups vulnerable to trace element toxicity, 1) children aged 1 to 3 years and 2) pregnant women aged 19 to 30 years (including those who plan to become pregnant and those who are breastfeeding). Recommended Dietary Allowances (RDA), Adequate Intake (AI) and Tolerable Upper Intake Level (UL) values were calculated for a weekly consumption of 150 g (pregnant women) or 75 g (children) of adult seal muscle or liver, as well as winter YY (< 6 weeks old) muscle or liver. Scenarios were not calculated for heart and kidney tissues because essential elements concentrations in these tissues were intermediate between muscle and liver tissues.

## 3. RESULTS

### 3.1 Trace element concentrations

Mean concentrations of trace elements in grey seal tissues are shown as a function of age category, sex, sample site and tissue type with ANOVA and post-hoc results in the Supplementary Material (Tables S5 to S13). Many of the trace elements measured in this study were not detected in liver (Sb, As, Be, B, Cr, Co, Pb, Ni, Tl, Sn) or muscle tissues (same elements as liver, plus Cd, Mn, Mo) for the first group of seal samples (Group 1). For the second group of samples (Group 2), detected elements were the same as for Group 1 plus As (muscle, heart, kidney), and Cd, Co and Pb (kidney) which were also detected because of lower analytical LOD for Group 2 analyses (Table S1). Results for Hg and MeHg are detailed separately in Section 3.2.

#### 3.1.1 Age differences

For muscle and liver tissues, the concentration of many of the detected trace elements varied significantly with seal age category (Table 1). For muscle tissues, Fe, Mg, and Se concentrations were significantly lower in winter YY (< 6 weeks old) when compared to spring YY (5 - 6 months old), juveniles (1 - 4 years) and adults (> 4 years). Muscle Cu concentrations were about 20% higher in winter and spring YY when compared to juveniles and adults. No trend was observed for Zn in muscle tissue (Table S5). For liver tissues, concentrations of Cd, Cr, Mn, Mo and Se were also lower in winter YY (< 6 weeks old) when compared to juveniles and adults. Conversely, concentrations of 3 essential elements (Fe, Mg, Zn) were higher in the livers of winter YY compared to juveniles and adults. Hepatic concentrations of Pb and Cu did not vary significantly with age category (Table S6). Liver samples were not collected from spring YY (< 6 months old) in Group 2 and we could not thoroughly test variation in kidney and heart concentrations by age category because only two categories were represented in Group 2 (spring YY and juveniles). For the majority of elements, no significant difference in hepatic or renal concentration were found between spring YY and juveniles, with the exception of Cd and Se which were significantly lower (24% and 83% respectively) in the kidneys of spring YY compared to juvenile seals (Table S7).

**Table 1.** Summary table for sample size (n), geometric means, geometric 95% confidence intervals (95% CI: lower limit, upper limit), and range (minimum - maximum) for trace element concentrations in µg/g w.w. measured in grey seals (*Halichoerus grypus*). Concentrations shown by age category and tissue type. Age categories were grouped into seals older than 5 months (adult, juvenile, spring YY seals, ∼ 5 months to 29 years) and seals younger than 6 weeks (winter YY). Concentrations are reported for analytes where more than 60% of samples were detected and concentrations under the LOD were replaced with half the LOD.

| Muscle |  | n | Cd | Cr | Co | Cu | Fe | Pb | Mg | Mn | Mo | Se | Zn |
| --- | --- | --- | --- | --- | --- | --- | --- | --- | --- | --- | --- | --- | --- |
| > 5 months | Mean | 85 | n.d. | n.d. | n.d. | 1.15 | 173 | n.d. | 262 | n.d. | n.d. | 0.385 | 36.5 |
|  | 95%CI |  | n.d. | n.d. | n.d. | (1.10, 1.20) | (164, 183) | n.d. | (254, 270) | n.d. | n.d. | (0.369, 0.401) | (34.4, 38.6) |
|  | Range |  | n.d. | n.d. | n.d. | 0.75 - 1.80 | 87 - 350 | < LOD - 0.48 | 180 - 380 | n.d. | n.d. | 0.24 - 0.66 | 18 - 92 |
| < 6 weeks | Mean | 34 | n.d. | n.d. | n.d. | 1.29 | 66.8 | n.d. | 222 | n.d. | n.d. | 0.340 | 39.9 |
|  | 95%CI |  | n.d. | n.d. | n.d. | (1.21, 1.37) | (61.6, 72.4) | n.d. | (210, 235) | n.d. | n.d. | (0.312, 0.370) | (37.6, 42.3) |
|  | Range |  | n.d. | n.d. | n.d. | 0.82 - 1.80 | 46 - 150 | < LOD - 0.20 | 160 - 320 | n.d. | n.d. | 0.20 - 0.57 | 27 - 59 |
| Total | Mean | 119 | n.d. | n.d. | n.d. | 1.19 | 132 | n.d. | 250 | n.d. | n.d. | 0.371 | 37.4 |
|  | 95%CI |  | n.d. | n.d. | n.d. | (1.15, 1.23) | (120, 144) | n.d. | (243, 258) | n.d. | n.d. | (0.357, 0.386) | (35.8, 39.1) |
|  | Range |  | n.d. | n.d. | n.d. | 0.75 - 1.80 | 46 - 350 | < LOD - 0.48 | 160 - 380 | n.d. | n.d. | 0.20 - 0.66 | 18 - 92 |
| Liver |  |  |  |  |  |  |  |  |  |  |  |  |  |
| > 5 months | Mean | 54 | 1.07 | 0.559 | 0.0128 | 31.4 | 294 | n.d. | 192 | 4.12 | 0.530 | 12.7 | 65.7 |
|  | 95%CI |  | (0.881, 1.30) | (0.459, 0.680) | (0.0114, 0.0143) | (26.8, 36.7) | (253, 340) | n.d. | (187, 197) | (3.86, 4.39) | (0.495, 0.567) | (9.87, 16.3) | (59.4, 72.7) |
|  | Range |  | 0.28 - 51 | 0.25 - 2.6 | <LOD - 0.027 | 7.2 - 83 | 65 - 1300 | < LOD - 0.048 | 150 - 230 | 2.4 - 6.4 | 0.29 - 0.87 | 2.2 - 110 | 20 - 150 |
| < 6 weeks | Mean | 35 | n.d. | 0.410 | n.d. | 25.5 | 619 | n.d. | 220 | 3.16 | 0.393 | 0.727 | 111 |
|  | 95%CI |  | n.d. | (0.335, 0.502) | n.d. | (22.2, 29.2) | (502, 763) | n.d. | (214, 227) | (3.02, 3.32) | (0.368, 0.419) | (0.670, 0.789) | (103, 119) |
|  | Range |  | n.d. | <LD - 1.1 | <LOD - 0.011 | 13 - 57 | 160 - 1700 | < LOD - 0.026 | 180 - 260 | 2.4 - 4.6 | 0.26 - 0.61 | 0.47 - 1.2 | 73 - 160 |
| Total | Mean | 89 | 0.245 | 0.495 | n.d. | 28.90 | 394 | n.d. | 203 | 3.71 | 0.471 | 4.12 | 80.7 |
|  | 95%CI |  | (0.163, 0.367) | (0.43, 0.57) | n.d. | (25.9, 32.3) | (342, 454) | n.d. | (198, 207) | (3.53, 3.91) | (0.445, 0.499) | (2.95, 5.76) | (74.1, 87.8) |
|  | Range |  | < LOD - 51 | <LD - 2.6 | < LOD - 0.027 | 7.2 - 83 | 65 - 1700 | < LOD - 0.048 | 150 - 260 | 2.4 - 6.4 | 0.26 - 0.87 | 0.47 - 110 | 20 - 160 |
| Kidney |  |  |  |  |  |  |  |  |  |  |  |  |  |
| > 5 months | Mean | 31 | 0.986 | n.d. | 0.0107 | 2.89 | 77.3 | 0.0166 | 158 | 0.903 | 0.117 | 2.08 | 20.3 |
|  | 95%CI |  | (0.734, 1.33) | n.d. | (0.00961, 0.0119) | (2.77, 3.03) | (71.3, 83.7) | (0.0111, 0.0248) | (153, 164) | (0.852, 0.958) | (0.108, 0.127) | (1.95, 2.21) | 19.7, 21.0) |
|  | Range |  | 0.27 - 4.3 | n.d. | < LOD - 0.026 | 2.1 - 3.5 | 45 - 110 | < LOD - 0.22 | 130 - 190 | 0.58 - 1.2 | 0.074 - 0.26 | 1.3 - 2.7 | 17 - 24 |
| Heart |  |  |  |  |  |  |  |  |  |  |  |  |  |
| > 5 months | Mean | 31 | n.d. | n.d. | n.d. | 2.79 | 87.5 | n.d. | 220 | 0.369 | 0.0323 | 0.439 | 26.5 |
|  | 95%CI |  | n.d. | n.d. | n.d. | (2.65, 2.93) | (80.2, 95.3) | n.d. | (212, 227) | (0.346, 0.394) | (0.0302, 0.0346) | (0.418, 0.460) | (25.5, 27.6) |
|  | Range |  | n.d. | n.d. | n.d. | 1.9 - 3.4 | 63 - 220 | < LOD - 0.26 | 170 - 260 | 0.21 - 0.5 | 0.023 - 0.045 | 0.32 - 0.64 | 18 - 24 |

The exact age of a subset of seals (n = 54) was determined by the number of tooth cementum layers in the canines. For 23 adult seals (19 males, 4 females) from Group 1, seal age ranged from 11 to 29 years old. All seals from Group 2 (n = 31) were analyzed for exact age and there were 11 juveniles in this group (aged 1.5 to 3.5 years) and 20 spring YY (aged 5 or 6 months). No significant associations were observed between exact age (years) and trace element concentrations (Cd, Cr, Cu, Fe, Mg, Mn, Mo, Se) in muscle (n = 54) or liver (n = 23) tissues for this subset of seals (Fig. 2), with one exception for Zn muscle concentrations, however Zn trends with exact age were not significant when testing adult seals only (Pearson’s r correlations, Table S14).

**Figure 2.**
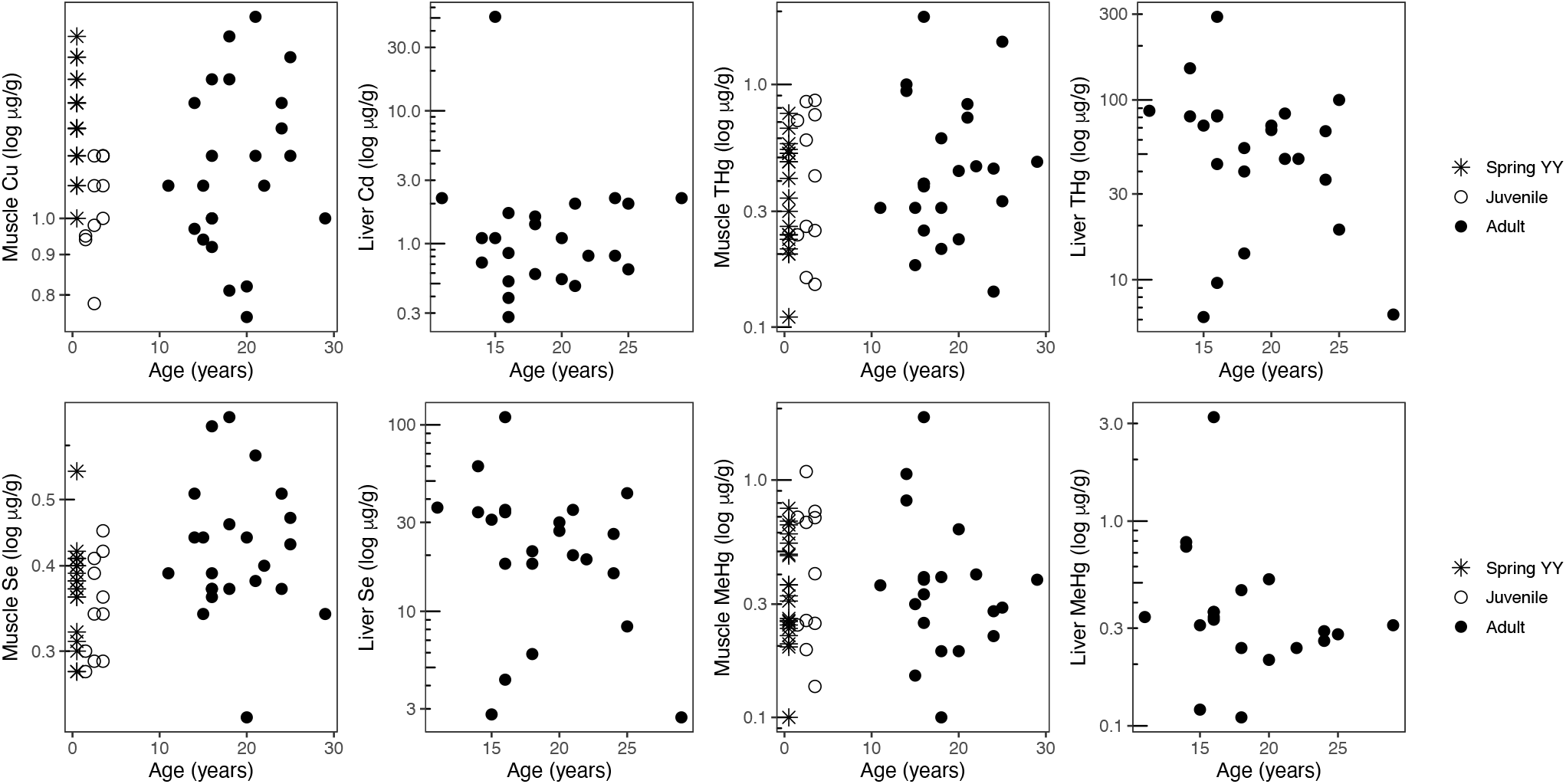
Correlations between exact age of grey seals (years) and selected trace element concentrations in log µg/g w.w. for muscle (THg, MeHg) and liver (THg, MeHg, Cd, Se) tissues. Exact age was determined for a subset of 54 seals, including 23 adults, 11 juveniles, 20 Spring YY. Sample size (n) varied for different elements: n = 20-54. No significant correlations (Pearson’s r, p > 0.05).

#### 3.1.2 Sex and site-specific differences

Trace element concentrations were analyzed as a function of sex and sampling site for muscle and liver tissues from a subset of juvenile and adult seals from Group 1 (muscle : n = 65, liver : n = 54). This subset was selected to reduce variation due to seal age category and because seals from Group 2 were all collected from the same site (Island 1). Hepatic Mn and Zn were more concentrated in male seals but no differences were observed for muscle tissues, except for mercury (see section 3.2.1) (Table S10-S11). No other significant differences were observed between the sexes. For sample site, trace elements concentrations were highly variable between sites. Higher concentrations of some elements were observed in muscle (Cu) and liver (Cr, Fe, Se, Zn) tissues from seals harvested at Island 4 (Table S12-S13).

#### 3.1.3 Tissue-specific differences

More elements were detected in liver (8) compared to muscle (5) tissues across all age categories, indicating that several elements (Cd, Mn, Mo) were more concentrated in the liver. For elements detected in both tissues (Cu, Co, Fe, Mg, Se, Zn), average hepatic concentrations were 1.8 to 35 times higher compared to muscle tissue (adults only, Student’s t-test, p ≤ 0.001), with the exception of Mg which was more concentrated in the muscle (t = 12.37, p ≤ 0.001). For the muscle, heart and kidney samples harvested from spring YY and juvenile seals (Group 2 samples), several elements were only detected in the kidneys (Cd, Co, Pb) (Table S1). As and Cu concentrations were 2-fold higher in hearts and kidneys compared to muscle samples, whereas the reverse was true for Fe. For Pb, the majority of muscle, liver and heart concentrations were below the LOD (< 0.01 µg/g for Group 1; < 0.005 µg/g for Group 2). When detected, Pb concentrations ranged from 0.006 - 0.48 µg/g in muscle and from 0.01 - 0.05 µg/g in liver. Renal Pb was detected in the majority (94%) of samples and ranged from 0.005 - 0.22 µg/g (Group 2). There were 8 outliers for Pb in seal muscle and 3 outliers in heart tissues (Fig. 3)

**Figure 3.**
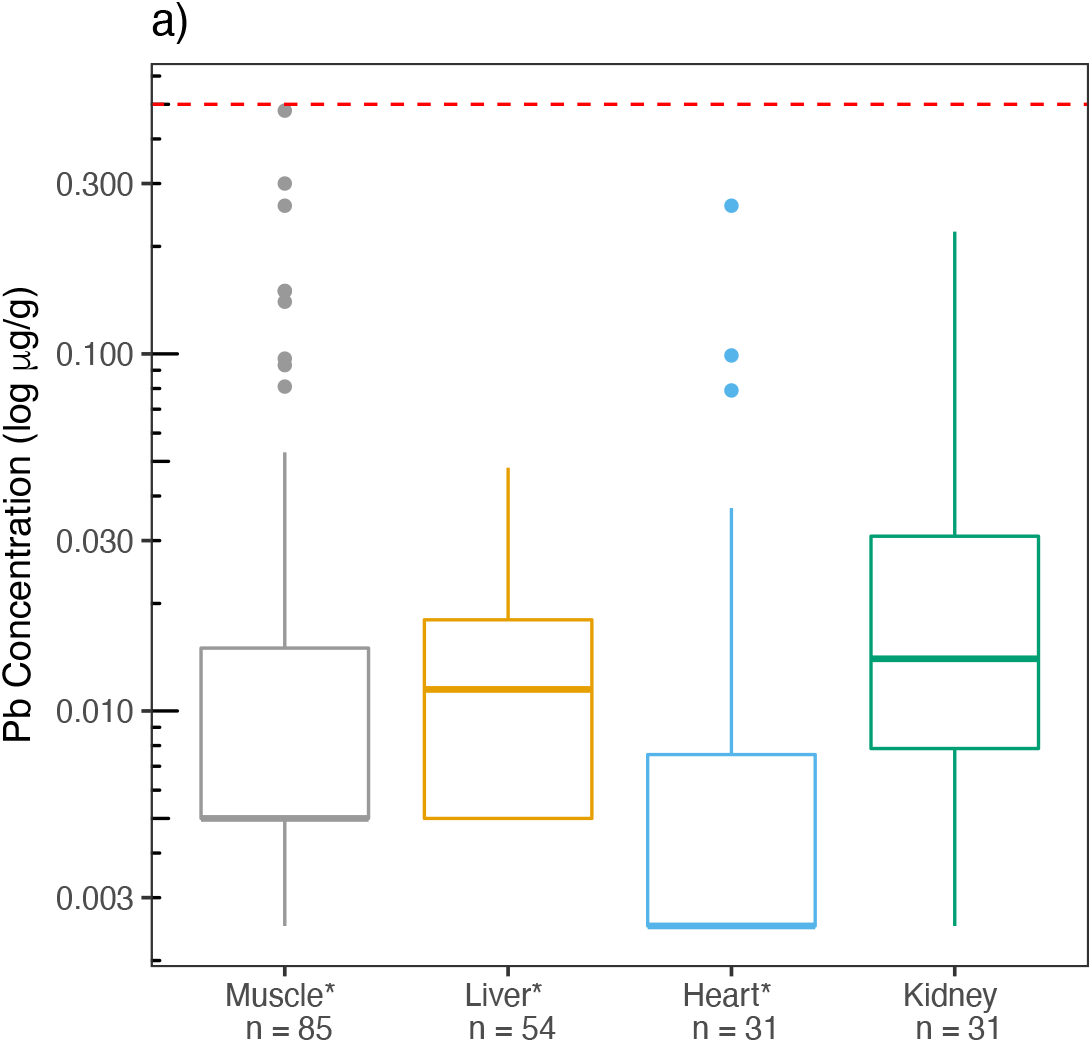
Concentrations in log10 ug/g w.w. of lead (Pb) in grey seals by tissue type. Boxplot values are medians (± 1st and 3rd quartiles) and points outside whiskers are outliers (outliers: muscle, n = 8, heart, n = 3). The asterisk (*) indicates that Pb was detected in less than 60% of muscle, liver and heart samples and values < LOD were replaced with half the LOD. Dotted line shows Canadian reference value for Pb of 3.5 µg/g w.w. in food products.

For the other elements (Mg, Mn, Hg, Mo, Se, Zn), concentrations varied significantly between muscle, heart and kidney tissues without showing a general trend. For example, Mg and Zn concentrations increased in the order kidney < heart < muscle, but in the opposite order for Mn. Since liver was not sampled for Group 2, hepatic concentrations from juvenile seals in Group 1 (n = 6) were compared with heart and kidney concentrations in juveniles from Group 2 (n = 11). The majority of trace elements (Cu, Fe, Mn, Hg, Se, Zn) were more concentrated in the liver than in hearts or kidneys for these juvenile seals, with the exception of Mg which increased in the order kidney < heart = liver < muscle (p ≤ 0.001). Renal Cd concentrations were significantly higher compared to liver samples for juvenile seals (2.44 µg/g versus 1.00 µg/g respectively).

### 3.2 Mercury and methylmercury

#### 3.2.1 Age-, sex- and site- specific differences

Similar to the other trace elements, THg and MeHg concentrations were significantly lower in winter YY (< 6 weeks old) compared to other age categories for both muscle and liver tissues (Table 2, Fig. 4, Table S8). No significant differences were observed for THg and MeHg in hearts and kidneys for spring YY and juvenile seals (with one exception, Table S9). No significant correlations were observed between exact age (years) and the concentrations of THg and MeHg in muscle and liver tissues (Fig. 2). However, significant differences were observed in THg and MeHg concentrations between the sexes. On average, THg and MeHg were more concentrated in the muscle tissues of adult and juvenile male seals. Average muscle concentrations were 0.39 µg THg/g and 0.43 µg MeHg/g for male seals and these concentrations were 1.6 (THg) and 1.7 (MeHg) times higher than in female seals (Table S10). No significant differences by sex were found for liver tissues. For sample site, THg concentrations were higher in muscle and liver tissues of seals collected on Island 4, when compared to Island 1 and 2 (p < 0.01, but no difference with Island 3). Among all samples, there were 4 adult seals (2 males and 2 females) with THg concentrations greater than 100 µg/g in the liver: 2 were from Island 4, 1 from Island 3, and 1 from Island 1. No significant differences were observed for MeHg concentrations in muscle or liver among sample sites (Tables S12-13).

**Table 2.** Summary table for sample size (n), geometric means, geometric 95% confidence intervals (95% CI: lower limit, upper limit), and range (minimum - maximum) for mercury concentrations in µg/g w.w. measured in grey seals (*Halichoerus grypus*). Concentrations shown by age category and tissue type. Age categories were grouped into seals older than 5 months (adult, juvenile, spring YY seals, ∼ 5 months to 29 years) and seals younger than 6 weeks (winter YY).

| **Muscle** |  | n | THg | n | MeHg | n | % MeHg |
| --- | --- | --- | --- | --- | --- | --- | --- |
| > 5 months | Mean | 85 | 0.341 | 51 | 0.353 | 51 | 85.3 |
|  | 95 % CI |  | (0.299, 0.389) |  | (0.298, 0.419) |  | (83.3, 87.4) |
|  | Range |  | 0.086 - 1.90 |  | 0.100 - 1.84 |  | 62.1 - 99.9 |
| < 6 weeks | Mean | 34 | 0.0430 | 12 | 0.0710 | 12 | 76.7 |
|  | 95 % CI |  | (0.0325, 0.0569) |  | (0.0525, 0.0961) |  | (70.7, 83.2) |
|  | Range |  | <LOD - 0.180 |  | 0.03 - 0.140 |  | 63.5 - 93.0 |
| Total | Mean | 119 | 0.189 | 63 | 0.260 | 63 | 83.6 |
|  | 95 % CI |  | (0.153, 0.233) |  | (0.210, 0.323) |  | (81.5, 85.8) |
|  | Range |  | <LOD - 1.90 |  | 0.03 - 1.84 |  | 62.1 - 99.9 |
| **Liver** |  |  |  |  |  |  |  |
| > 5 months | Mean | 54 | 28.7 | 26 | 0.345 | 26 | 0.795 |
|  | 95 % CI |  | (21.9, 37.6) |  | (0.268, 0.446) |  | (0.549, 1.15) |
|  | Range |  | 3.80 - 290 |  | 0.110 - 3.22 |  | 0.270 - 5.16 |
| < 6 weeks | Mean | 35 | 0.283 | 19 | 0.0720 | 19 | 24.2 |
|  | 95 % CI |  | (0.214, 0.374) |  | (0.0554, 0.0936) |  | (21.1, 27.8) |
|  | Range |  | 0.072 - 2.20 |  | 0.03 - 0.170 |  | 12.9 - 39.6 |
| Total | Mean | 89 | 4.67 |  | 0.178 |  | 3.36 |
|  | 95 % CI |  | (2.79, 7.82) | 45 | (0.133, 0.239) | 45 | (1.93, 5.86) |
|  | Range |  | 0.072 - 290 |  | 0.03 - 3.22 |  | 0.270 - 39.6 |
| **Kidney** |  |  |  |  |  |  |  |
| > 5 months | Mean | 31 | 0.678 | 31 | 0.266 | 31 | 33.4 |
|  | 95 % CI |  | (0.535, 0.858) |  | (0.211, 0.334) |  | (30.3, 36.9) |
|  | Range |  | 0.17 - 2.90 |  | 0.072 - 0.918 |  | 16.4 - 53.0 |
| **Heart** |  |  |  |  |  |  |  |
| > 5 months | Mean | 31 | 0.128 | 31 | 0.118 | 31 | 87.7 |
|  | 95 % CI |  | (0.103, 0.159) |  | (0.0942, 0.149) |  | (86.0, 89.4) |
|  | Range |  | 0.045 - 0.50 |  | 0.037 - 0.448 |  | 76.9 - 96.6 |

**Figure 4.**
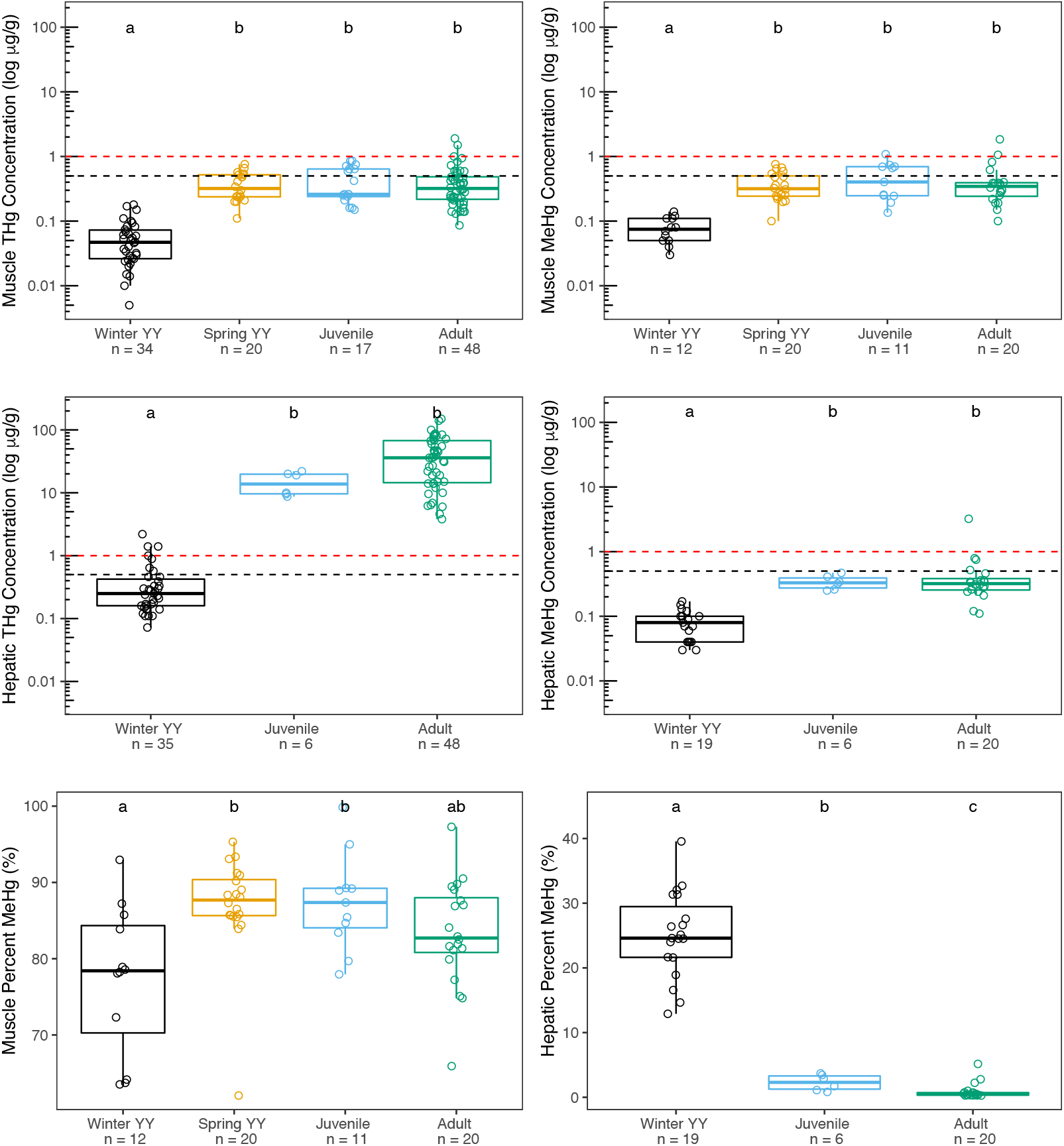
Concentrations in log10 ug/g w.w. of THg (left) and MeHg (right) in grey seal muscle and liver tissues shown by age category. Boxplot values are medians (± 1st and 3rd quartiles) and points outside whiskers are outliers. Dotted lines indicate Canadian reference values of 0.5 (black) and 1.0 (red) µg/g w.w. of THg. Letters show results of one-way ANOVAs with Post-Hoc Tukey HSD tests.

Two-factor ANOVA with post-hoc analyses (Tukey test) was used to estimate the relative importance of sex (male, female) and age (4 categories) on mercury concentration in seal tissues. Sample site could not be included as a factor in the multivariate model because of the unbalanced study design, i.e., all age categories were not sampled at each location. For THg in muscle tissues, both age category and sex were statistically significant (p < 0.01). The effect size of age category was 0.382, indicating that 38.2% of the variation of THg concentration in muscle tissues was explained by age category (F = 29.4, p < 0.001), more specifically by the difference in concentrations between winter YY and all other categories (with no difference between the other categories) (Tukey HSD, p < 0.001). Effect size for sex was 0.034, indicating that only 3.4% of the variation in THg in muscle was explained by sex (F = 7.89, p = 0.006), hence by the higher concentrations found in male seals (Tukey HSD, p < 0.001). For THg in liver tissues, only age category was significant (F = 123.2, p < 0.001) with an effect size of 0.651 or 65.1% explained by lower THg concentrations found in winter YY (< 6 weeks old). Models for MeHg concentration in muscle and liver tissues were very similar to models for THg. For MeHg in seal muscle, both variables were significant and the effect size for age category was 46.9% (F = 22.9, p < 0.001), which was well above the 3.8% effect size for sex (F = 5.56, p = 0.022). For hepatic MeHg, only age category was significant (F = 21.2, p < 0.001) with an effect size of 47.1%. None of the interaction terms from the multivariate models were significant (p > 0.05) indicating that there was no combined effect of age category and sex on THg or MeHg concentrations.

#### 3.2.2 Tissue-specific differences

Regardless of age category, hepatic THg concentrations were higher (31-fold on average) compared to muscle tissues in the same seal. As shown in Table 2, after grouping all age categories together (except winter YY to control for age-specific differences), THg concentrations increased in the following order: heart < muscle < kidney < liver, with an approximately 100-fold difference between heart and liver tissues (F = 521.3, p < 0.001). MeHg concentrations were much less variable among the different tissues (Table 2). Considering all age categories (except winter YY), MeHg concentrations were significantly lower in heart tissues compared to the other tissue types (F = 22.64, p < 0.001). THg and Se showed a very strong positive correlation in the liver (r = 0.97, p < 0.01) but less so in the muscle (r = 0.46, p < 0.01), heart (r = 0.36, p = 0.04) and kidney (r = 0.39, p = 0.03). MeHg and Se were likewise strongly correlated in the liver (r = 0.85, p < 0.01), less strongly so in the heart (r = 0.45, p = 0.01), and were not correlated in muscle (r = 0.10, p = 0.41) or kidney (r = 0.21, p = 0.26) tissues.

The percentage of total mercury as methylmercury (% MeHg) was calculated for different tissues by age category (Tables S8- S9). Across all age categories, % MeHg averaged 83% (range: 62 - 100%) in muscle tissues. The % MeHg was much lower in liver tissues (3.4 %) and more variable with seal age (range: 0.3 - 40 %). For heart tissues, the % MeHg was similar to muscle tissues and averaged 88 % (range: 77 - 97 %) and in the kidneys, average % MeHg was 33 % (range: 16 - 53 %). Percent MeHg also varied significantly with age category (Fig. 4). For muscle tissues, average % MeHg was significantly lower in winter YY (76 %) compared to spring YY and juvenile seals (87 %) (p = 0.0029), but was not different from adult seals whose % MeHg values were 83% (range 65 % - 97 %). On average, hepatic % MeHg was much higher in winter YY (24 %, range 21 - 28 %) compared to juveniles and adults whose % MeHg was very low (< 5 %). The % MeHg also decreased with age in the kidneys with an average of 38 % in spring YY to 26 % in juvenile seals (p < 0.001) (Tables S8-S9).

### 3.3 Comparison with Reference Values

#### 3.3.1 Reference values for essential trace elements

For essential trace elements detected in grey seal tissues in this study (Cr, Cu, Fe, Mg, Mn, Mo, Se, Zn), the Percent Dietary Reference Intakes (% DRI) were calculated for two vulnerable human population subgroups using four consumption scenarios (Table 3, Table S15). Recommended Dietary Allowances (RDA), Adequate Intake (AI) and Tolerable Upper Intake Level (UL) values were calculated for grey seal muscle and liver tissues from adult seals (> 4 years) and from winter young-of-the-year (winter YY, < 6 weeks) based on weekly consumption of 75 g of seal tissue for children 1-3 years old and 150 g of seal tissue per week for pregnant women 19 - 30 years old.

**Table 3.** Comparison of grey seal essential and non-essential trace elements concentrations to Canadian reference values. For essential elements, this table shows the percentage of Tolerable Upper Intake Level (% UL) based on consumption scenarios of 75 g of muscle or liver tissue weekly for children from 1 to 3 years. Levels are exceeded when % UL is above 100 %. For non-essential elements, this table shows the percentage of samples above Canadian reference values for food products (% > CRV) and values are exceeded when above 0 %.

| Essential elements : Percentage of Tolerable Upper Intake Level (% UL) based on consumption of 75 g of muscle or liver tissue weekly for children from 1 to 3 yr. |  |  |  |  |  | Non-essential elements : Percentage of grey seal samples above Canadian reference values (% > CRV) for food products. |  |  |  |  |  |
| --- | --- | --- | --- | --- | --- | --- | --- | --- | --- | --- | --- |
| Winter young-of-the-year grey seals (less than 6 weeks old) |  |  |  |  |  | Winter young-of-the-year grey seals (less than 6 weeks old) |  |  |  |  |  |
|  |  | Muscle |  | Liver |  |  |  | Muscle |  | Liver |  |
| Essential | UL | % UL | Intake | % UL | Intake | Non-essential | CRV | % > CRV | µg/g | % > CRV | µg/g |
| Copper (Cu) | 3 000 µg/day | 1.4 % | Low | 27 % | Low | Arsenic (As) | 3.5 µg/g | 0 % | As < 0.10 | 0 % | As < 0.10 |
| Iron (Fe) | 40 000 µg/day | 1.8 % | Low | 17 % | Low | Cadmium (Cd) | 1.5 µg/g* | 0 % | Cd < 0.05 | 0 % | Cd < 0.05 |
| Selenium (Se) | 150 µg/day | 4.0 % | Low | 8.7 % | Low | Lead (Pb) | 0.5 µg/g | 0 % | Pb < 0.05;<br>one outlier | 0 % | Pb < 0.03 |
| Zinc (Zn) | 12 000 µg/day | 6.1 % | Low | 17 % | Low | Total mercury (THg)<br>Methylmercury (MeHg) | 0.5 µg/g | 0 %<br>0 % | THg < 0.18<br>MeHg < 0.14 | 20 %<br>0 % | Hg total < 2.2<br>MeHg < 0.17 |
| Adult grey seals only |  |  |  |  |  | Adult, juvenile, and spring young-of-the-year grey seals |  |  |  |  |  |
|  |  | Muscle |  | Liver |  |  |  | Muscle |  | Liver |  |
| Essential | UL | % UL | Intake | % UL | Intake | Non-essential | CRV | % > CRV | µg/g | % > CRV | µg/g |
| Copper (Cu) | 3 000 µg/day | 1.2 % | Low | 32 % | Low | Arsenic (As) | 3.5 µg/g | 0 % | As < 0.28 | 0 % | As < 0.10 |
| Iron (Fe) | 40 000 µg/day | 4.4 % | Low | 8.0 % | Low | Cadmium (Cd) | 1.5 µg/g* | 0 % | Cd < 0.05 | 20 % | Cd < 51.0 |
| Selenium (Se) | 150 µg/day | 4.7 % | Low | 165 % | High** | Lead (Pb) | 0.5 µg/g | 0 % | Pb < 0.08<br>outliers | 0 % | Pb < 0.05 |
| Zinc (Zn) | 12 000 µg/day | 6.0 % | Low | 10 % | Low | Total mercury (THg)<br>Methylmercury (MeHg) | 0.5 µg/g | 29 %<br>27 % | THg < 1.90<br>MeHg < 1.84 | 100 %<br>15 % | Hg total < 290<br>MeHg < 3.22 |
|  |  |  |  |  |  |  |  | Heart |  | Kidney |  |
|  |  |  |  |  |  | Non-essential | CRV | % > CRV | µg/g | % > CRV | µg/g |
|  |  |  |  |  |  | Arsenic (As) | 3.5 µg/g | 0 % | As < 0.75 | 0 % | As < 0.63 |
|  |  |  |  |  |  | Cadmium (Cd) | 1.5 µg/g** | 0 % | Cd < 0.05 | 29 % | Cd < 4.3 |
|  |  |  |  |  |  | Lead (Pb) | 0.5 µg/g | 0 % | Pb < 0.01<br>outliers | 0 % | Pb < 0.22 |
|  |  |  |  |  |  | Total mercury (THg)<br>Methylmercury (MeHg) | 0.5 µg/g | 3.2 %<br>0 % | THg < 0.50<br>MeHg < 0.45 | 55 %<br>22 % | THg < 2.9<br>MeHg < 0.92 |

\*There is no Canadian reference value for Cd in food products, see section 3.3 for more details.
\*\* Excessive consumption of Fe or Se is unlikely to result in adverse health effects, see section 4.4 for more details.

Based on the % RDA values, essential trace elements concentrations in muscle tissues from seals of either age category would contribute less than 30% of the recommended intake values for both pregnant women and young children. This means that the consumption of a weekly portion of seal muscle harvested from either adult or YY seals would provide less than the minimum recommended intake of essential elements for vulnerable population subgroups. It therefore follows that, based on the Tolerable Upper Intake Level (UL) values, the weekly consumption of 150 g or 75 g of seal muscle respectively would not lead to excessive intake of any of these essential elements (Table 3, Table S15).

Based on the % RDA values, the consumption of a weekly portion of adult seal liver would be a good source of Cu (93 % for children; 63 % for pregnant women) but would provide more than the minimum recommended intake for Se (742 % for children; 495 % for pregnant women). Indeed, based on the UL values for scenarios with adult seal liver, a weekly serving would exceed the recommended maximum values for Se for children 1- 3 years old (165 %). Based on scenarios with winter YY liver, a weekly portion would be a good dietary source of Cu (80 % for children; 55 % for pregnant women) and Fe (95 % for children; 49 % for pregnant women) and would not lead to excessive intake of any of these essential elements (% UL < 100).

#### 3.3.2. Reference values for non-essential trace elements

Several non-essential trace elements were detected in grey seal tissues in this study, including As (muscle, heart, kidney), Cd (liver, kidney), Pb (kidney), and THg and MeHg (muscle, liver, heart, kidney). When detected, the concentrations of As were less than 1 µg/g (mean < 0.3 µg/g, range 0.05 - 0.75), and well below the Canadian reference value of 3.5 µg/g for As in food. When detected, all Pb concentrations were below the Canadian reference value of 0.05 µg/g for fish protein. Cd was not detected in muscle or heart tissues, but was detected in 61% of liver samples and 100% of kidney samples. Although there is no Canadian reference value for Cd in food, other studies on cervids have used an intake limit of 1.0 or 1.5 µg/g for Cd following guidelines from the World Health Organization (Robillard et al., 2002; Stansley et al., 1991). None of the liver samples from winter YY exceeded the 1.5 µg Cd/g value. For the other age categories, 20% of liver (n = 11/54) and 29% of kidney samples (n = 9/31, Group 2 only) exceeded this 1.5 µg Cd/g intake limit (Table 3).

The reference value for THg in Canada is either 0.5 µg/g for most commercially sold fish products or 1.0 µg/g for a few species, including fresh/frozen tuna, shark, swordfish, marlin, orange roughy and escolar (Canada, 2007). In the present study, as shown in Fig. 4, none of the muscle samples from winter YY exceeded these recommended maximum concentrations for THg or for MeHg. For livers from winter YY, 20 % and 11 % of samples exceeded these 0.5 and 1.0 µg/g values respectively based on THg concentrations, but none (0 %) exceeded these values based on MeHg concentrations. For the other age categories all together, 29% of muscle samples, 100 % of liver samples, 55 % of kidney samples, and 3.2 % of the heart samples exceeded 0.5 µg/g of THg. However, these proportions were lower when compared to MeHg, specifically 27 % for muscle, 15 % for liver, 22 % for kidney, and 0 % for heart tissues. When comparing to the 1.0 µg THg/g reference value, percentages were much lower, namely 5.9 % for muscle, 3.8 % for liver, and 0 % for kidney and heart tissues. Although samples from adult male seals more frequently exceeded these reference values for THg and MeHg in the liver, this trend may have been influenced by the small number of adult females (n=11) compared to males (n=37). As presented earlier, Hg concentration was found to be slightly higher for male seals, but did not show a significant increase with age after 5 months. Indeed, for MeHg in muscle samples, there were almost equal numbers of spring YY, juveniles and adults sample that exceeded the 0.5 µg/g value (36%, 36% and 28% respectively).

## 4. DISCUSSION

### 4.1 Age, sex and site-specific differences in trace element concentrations

In this study, we did not observe progressive bioaccumulation of trace elements with age in either muscle or liver tissues. Contrary to other studies, trace element (including mercury) bioaccumulation in grey seals in the present study showed two distinct groups based on age category; winter young-of-the-year (winter YY, < 6 weeks old) and all other age categories together (Figs. 2, 4). Although there are several routes of exposure for trace elements, the primary route of uptake in seals is through food (Das et al., 2002), hence these results likely reflect a change in diet following weaning. The lack of significant differences in most trace elements concentrations among spring young-of-the-year (spring YY, 5-6 months), juveniles (1 - 4 years) and adults (11 - 29 years) suggest a similar diet after weaning for a broad range of ages of grey seals in the GSL. Indeed, a previous study from the same region showed no significant difference in grey seals’ diet as a function of age with seals from 1 - 33 years old (Hammill et al., 2014). Although many studies on marine mammals show gradual and continuous bioaccumulation for Cd, Se, THg, and MeHg with age (Marino et al., 2011; Dehn et al., 2005; Bustamante et al., 2004; Dietz et al., 1996), this is not universally observed and lack of such trends may be due to species- or tissue-specific differences or the influence of study design (sample size, age range of seals, geographic area, or concentration range) (Lahaye et al., 2006; Fant et al., 2001). The lack of association of trace element concentrations with exact age (years) in this study also suggests that dietary uptake of potentially toxic elements (Cd, THg, MeHg) remains in equilibrium with physiological detoxification processes (Dehn et al., 2005).

We could not thoroughly test age-dependent bioaccumulation in the hearts and kidneys as these tissues were only sampled for two age categories. These tissues would need to be sampled from all age classes to better understand age-dependent trends. However, renal Cd and Se concentrations increased significantly from spring YY (5 - 6 months old) to juvenile seals (1 - 4 years old) likely indicating a progressive increase with age for these two elements.

In addition, and in contrast to Cd, Se, THg and MeHg, some essential elements were higher in the livers (Fe, Mg, Zn) and muscles (Cu) of winter YY (< 6 weeks old) when compared to juvenile and adult seals. Previous studies have also observed higher concentrations of Cu and Zn in young marine mammals (Das et al., 2002; Watanabe et al., 1998) which can be explained by an increase in essential elements during development for tissue differentiation and growth, or possibly by the limited excretion mechanisms of the fetus (Dehn et al., 2005; Wagemann et al., 1998).

Sex-specific differences in trace element bioaccumulation were not very pronounced in the present study. A few elements were ∼ 30 - 70% more concentrated in the liver (Mn, Zn) or muscle (THg, MeHg) tissues of male seals. These observed differences between the sexes could be due to size dimorphism, as male grey seals can weight up to 290 kg and females usually do not exceed 190 kg (Bustamante et al., 2004), or to physiological or metabolic dimorphism (Caurant et al., 1994; Robinson et al., 2012). Sex-related differences could also be due to differences in foraging behaviour between the sexes. Male grey seals in the GSL have been shown to eat a greater proportion of higher trophic-level fish, such as cod and hake, which likely have higher mercury concentrations than lower trophic-level species, like sandlance and herring, which are consumed in greater proportions by female seals (Hammill et al., 2014).

Previous studies on grey seals in the Faroe Island, the Baltic Sea, and from Sable Island (part of the same population as the present study) have found that female seals had higher levels of Cd in liver and kidney (Bustamante et al., 2004; Nyman et al., 2002) but this trend was not observed in the present study. One outlier female seal had extremely high levels of Cd (51 *µg/g*) but all other individuals had < 3.0 ug Cd/g. Finally, it was difficult to draw any firm conclusions about the influence of sample site on bioaccumulation in the present study because of the study design. Future studies may wish to examine the influence of harvest location on bioaccumulation in more detail, however it is important to note that grey seals from the GSL and the Northwest Atlantic are considered a single population and sampling sites do not represent distinct populations (Government of Canada, 2016b).

### 4.2 Tissue-specific differences in trace element concentrations

Tissue-specific bioaccumulation patterns have been observed in many previous studies on marine mammals where most trace elements concentrate to the liver and Cd concentrates to the kidney (AMAP, 1998; Nyman et al., 2002; Bustamante et al., 2004). Our results for grey seals mirrored this trend, with many elements (Cu, Co, Fe, Mn, Mo, THg, Se, Zn) more concentrated in the liver, and Cd highest in the kidney. Liver THg concentrations in the present study were 31 times higher on average than those in muscle tissues from the same seal, which is close to trends observed in ringed seals in northern Canada (Braune et al., 2015). Trace elements concentrate in the liver due to this organ’s role in the homeostatic regulation of essential trace elements and the sequestration of non-essential and toxic trace elements (Hansen et al., 2016), whereas Cd tends to accumulate in the kidneys due to the presence of metal-binding proteins (Das et al. 2002).

Pb concentrations were mostly below detection limits in muscle, liver and heart tissues in this study and these low levels are consistent with previous studies on marine mammals (AMAP, 1998; Fant et al., 2001; Nyman et al., 2002). Outliers for Pb in seal muscle (n = 8) and heart (n = 3) ranged from 0.08 to 0.48 µg/g in the present study (Fig. 3) and may indicate tissue contamination from lead bullets, as observed in a previous study on moose and deer (Fachehoun et al., 2015). However, harvesters in this study targeted the heads of the animals and outliers were found in samples taken from far inside the abdomen to avoid bullet fragments. One young seal (winter YY, < 6 weeks) hunted by hakapik also had a higher than average muscle Pb concentration (0.02 µg Pb/g). Given all of this, high Pb concentrations in some individuals may have been related to the use of lead bullets or to other natural or anthropogenic (e.g. lead fishing weights) sources of exposure.

Focusing on mercury, adult grey seal livers from had the highest THg concentrations (5 - 450 µg/g) of any tissue but had very low proportions of mercury in the form of methylmercury (range: 0.27 - 5.2 % MeHg). Muscle and heart tissues had much lower concentrations of THg (0.05 - 2.3 µg/g) but had high % MeHg (∼ 80%). Kidney samples were intermediate for both THg and % MeHg. Our results are similar to those of other studies on seals in Canada which show low proportions of MeHg in the liver (1-12 % MeHg in Braune et al., 2015) and higher proportions in seal muscle tissues (81 % in Dehn et al., 2005). The negative correlation between age category and % MeHg in seal livers in this study is also consistent with previous studies (cited in Ewald et al., 2019). These results, as well as the strong positive correlation between Hg and Se in liver tissues, likely reflect detoxification mechanisms, where mercury accumulates as stable and insoluble crystals of mercury selenide (HgSe) in the liver to protect marine mammals against exposure to MeHg (Ewald et al., 2019; Ikemoto et al., 2004). Moreover, based on toxicokinetic modeling for ringed seals in Ewald et al. (2019), cellular composition changes over time and an increase in thiol-containing compounds in the liver could also lead to increased partitioning of MeHg into inorganic Hg after about 4 - 5 months of age.

### 4.3 Geographical and inter-species comparison

Trace elements concentrations were in the same order of magnitude as those reported for other marine mammals, namely lower tissue concentrations (< 1.0 µg/g w.w.) for As, Cr, Co, Pb, Mo, and Ni and higher concentrations (∼ 1.0 - 1000 µg/g w.w.) for Cd, Cu, Fe, Hg, Mg, Mn, Se and Zn in some tissues (AMAP, 1998; Bustamante et al., 2004; Frank et al., 1992). In the following sections, all comparisons between studies were done using values for adult seals only to reduce variation due to different age ranges. Hepatic concentrations of Cu and Zn in the present study were similar to levels in other pinnipeds and grey seals from other regions (Bustamante et al., 2004; Das et al., 2002). Although Fe, Mg and Mn concentrations are not frequently reported in other studies, hepatic Fe concentrations for adult seals in the present study (range: 65 - 1300 µg/g) seem to be higher than previous reported values (range: 154 - 665 µg/g for grey seals from Sweden in Frank et al., 1992). However, the interpretation of results for Fe can be challenging because organs like the liver may be congested with blood, which contains large and variable quantities of Fe (Bratton et al., 2002).

Concentrations of As in this study (range: 0.05 - 0.75 µg/g) were on the low end of recorded values for other marine mammals (Kubota et al., 2001). Higher concentrations of As are often found in the blubber, especially for marine mammals feeding on cephalopods and crustaceans (Ebisuda et al., 2002), however this tissue type was not sampled in the present study. Low Pb levels (< 0.50 µg/g, including outliers) are consistent with previous studies on seals where concentrations > 1 µg Pb/g were considered high and attributed to industrial pollution (O’Shea, 1999) or sample contamination (Nyman et al., 2002).

In the present study, THg concentration ranges in adult grey seal muscle (0.086 - 1.90 µg/g) were lower than ranges for grey seals from the Faroe Islands sampled in 1993 - 1995 (0.13-4.61 µg/g, in Bustamante et al., 2004) and similar to ranges from Sable Island sampled in 1996 - 1998 (0.4 - 1.6 µg/g in Nyman et al., 2002). The mean calculated for adult seals in the present study was 0.34 µg THg/g muscle tissue, which is in the same range as means for ringed seals from across the Canadian Arctic (0.10 - 0.69 µg THg/g muscle tissue in Houde et al. (2020)) and on the low end of averages reported in other studies of Arctic ringed seals (Brown et al., 2016; Gaden et al., 2009). THg concentrations in grey seal liver are discussed in the next paragraph. Most studies on pinnipeds measure only the THg concentrations and do not measure Hg speciation, although MeHg is a better indicator of toxic effects. To the best of our knowledge, no other study has measured MeHg concentrations in grey seal organs (except for blood and milk in Grajewska et al., 2019). In the present study, MeHg concentrations for adult grey seals ranged from 0.10 - 1.84 µg/g in muscle and 0.11 - 3.00 µg/g in liver tissues which is similar to ranges for Arctic ringed seal muscle (0.13 - 1.09 µg/g) but lower than ranges for liver (2.50 - 89.9 µg/g) (Lemire et al., 2015).

Previous studies on grey seals have reported high renal Cd and hepatic Hg and Se concentrations when compared to other seal species, including ringed and harbour seals (Bustamante et al., 2004; Frank et al., 1992; Law et al., 1992; Nyman et al., 2002). Grey seals from the Faroe Islands had renal Cd and hepatic Hg and Se levels among the highest reported in the literature for temperate regions far from industrial contamination (Bustamante et al., 2004). Hepatic Hg concentrations for adult seals in the present study (3.8 - 290 µg/g) were similar to levels in Faroe Island grey seals (1.1 - 238 µg/g) and slightly lower than grey seals from a previous study of the geographically proximate Sable Island (15 - 348 µg/g in Nyman et al., 2002). These Hg concentrations in grey seal livers are higher than those for ringed seals from Svalbard and the eastern Canadian Arctic, but similar to hepatic Hg levels for ringed seals in the western Arctic (Fant et al., 2001; Wagemann et al., 1996). Although these studies in the Canadian Arctic are from 20 years ago, a recent study shows only very limited declines in Hg concentrations in Arctic ringed seals over the last five decades, despite declines in atmospheric mercury concentrations (Houde et al., 2020). Recent studies in the Canadian Arctic report average hepatic THg concentrations of 6.1 to 70.4 µg/g (Brown et al., 2016) and 1.8 to 27.8 µg/g (Houde et al., 2020) which overlap with averages calculated for juvenile and adult seals in the present study (13.8 and 31.5 µg THg/g of liver tissue, respectively).

In the present study, renal Cd ranges (0.27 - 4.3 µg/g) were lower than those measured in grey seals from the Faroe Islands (0.39 - 155 µg/g) and the Baltic sea (3.0 - 20 µg/g). Renal Cd was also slightly lower than concentrations measured 20 years ago from the same grey seal population as the present study (Sable Island, range: 1.4 - 7.6 µg/g in (Nyman et al., 2002). Renal Cd in the present study was also lower compared with seals and other marine mammals from the North American Arctic where Cd bioaccumulation is known to be higher (> 10 µg/g) in the liver and kidney, likely related to consumption of marine invertebrates such as amphipods (Brunborg et al., 2006; AMAP, 1998; Dietz et al., 1996). For example, average renal Cd ranged from 1.97 to 21.5 µg/g for ringed seal populations in Brown et al. (2016) compared to an average of 1.08 µg/g for adults in the present study. Hepatic Se ranges in the present study (2.2 - 110 µg/g for adult seals) were similar to those reported for grey seals from the Faroe Island (1.2 - 99 µg/g) and Sable Island (9.3 - 83 µg/g). Average hepatic Se concentrations in the present study (13.87 µg/g) were also very close to average concentrations from populations of ringed seals in the Canadian Arctic, which ranged from 3.49 to 14.7 µg/g in Houde et al. (2020).

### 4.4 Comparison with reference values

For essential elements, comparison to reference values based on consumption scenarios indicated that all elements were below the Recommended Dietary Allowances (% RDA < 100 %) for seal muscle tissues. This indicates that a weekly portion of 75 g (children 1 - 3 years) or 150 g (pregnant women 19 - 30 years) of either winter YY or adult seal meat does not pose a risk of excessive intake of essential elements for these vulnerable human population subgroups. The concentrations of many elements were higher in the liver, and therefore scenarios showed that a weekly portion of winter YY liver would be an excellent source of Cu and Fe without exceeding maximum tolerable intakes (UL) for these elements. A weekly portion of adult seal liver would also be an excellent source of Cu but would exceeded the maximum tolerable intakes (UL) for Se among children 1- 3 years old. For intakes above the UL, the potential risk of adverse effects to an individual increases but we cannot accurately estimate the proportion of the population who would experience adverse effects at any specific intake above the UL (Canada, 2005b).

However, it is important to note that Se accumulates in marine mammal livers as stable crystals of inorganic Se and Hg which are poorly assimilated by the intestine when ingested (Ikemoto et al., 2004). It is therefore unlikely that concentrations of Se in adult grey seal livers would result in adverse health effects related to nutrient overconsumption. We chose to use these consumption scenarios as they reflect a reasonable estimate of the average consumption of seal meat or liver in the Magdalen Islands, which is on a weekly, monthly, or more infrequent basis. Seal liver is also often sold or served in *pâte* where it is mixed with pork liver. Elsewhere in Quebec, seal products remain a delicacy served primarily in gourmet restaurants, although sport hunting for seal is gaining in popularity in Eastern Quebec.

For non-essential elements, the consumption of grey seal meat or tissues should not pose a risk for either As or Pb (despite some possible contamination by Pb in seal tissues). There may, however, be possible health risks associated with the moderately elevated hepatic and renal Cd concentrations in grey seals older than 5 months (Tables 3, S15). Importantly, as this is the age group primarily targeted during commercial hunts, none of the muscle samples from winter YY seals (< 6 weeks old) exceeded the recommended maximum concentrations for THg and MeHg. Moreover, although a few liver samples from winter YY exceeded maximum concentrations for THg, none of the liver samples exceeded the maximum concentrations when considering MeHg. For the other age classes (i.e. grey seals that have started feeding at sea), the majority of muscle (71%) and heart (97%) samples were below the recommended value of 0.5 µg/g for both THg and MeHg, but the majority of liver (100%) and kidney (55%) samples exceeded this maximum concentration for THg. However, Hg speciation analyses indicate that the percentage of MeHg in liver and kidney is much lower than in muscle and that, in the end, only 15% of the liver and 22% of the kidney samples have MeHg values above the recommended value of 0.5 µg/g, or only 3.8 % and 0 % respectively when considering the 1.0 µg/g recommended value for MeHg.

### 5. CONCLUSIONS

This study provides an exhaustive dataset on trace element bioaccumulation for grey seals in the Gulf of Saint Lawrence (GSL). Although we did not observe progressive age-dependent bioaccumulation, the results of this study indicate that seal age does play a major role in trace element concentration, most likely according to feeding behaviour. Indeed, our analyses showed two distinct groups, the winter young-of-the-year (< 6 weeks old) and all older seals grouped together (∼ 5 months - 29 years). After the start of feeding at sea, age did not appear to have a marked influence on trace element concentrations (including total mercury and methylmercury) likely due to similar diets over a broad range of seal ages.

Compared to other studies on pinnipeds, grey seals from the GSL had lower renal Cd concentrations and relatively high hepatic THg and Se concentrations. Despite lower Cd concentrations than in other seal populations, moderately elevated Cd levels in the liver and kidney of grey seals older 5 months and older suggest the need for continued vigilance regarding potential health risks. Grey seal tissues were well below the Canadian reference values for As and Pb, however some outlier concentrations for Pb in muscle and heart tissues raise the importance of promoting non-toxic ammunition for seal hunting. From a public health perspective, our results also highlight the importance measuring MeHg instead of total Hg for developing dietary recommendations for marine mammal consumption, such as grey seals from the GSL. Although the majority of liver and kidney samples from grey seals feeding at sea were above the recommended maximum values for THg in Canada, less than a quarter of samples exceeded recommended values based on the MeHg concentrations. Overall, our study indicates that, for winter young-of-the-year (< 6 weeks old), consumption of liver could be a good dietary source of essential elements (primarily Cu and Fe), and consumption of winter young-of-the-year muscle and liver can be done without exceeding maximum recommended concentrations for most trace elements. Ongoing discussions with regional public health professionals will help develop dietary recommendations for the consumption of the muscle, heart, kidney or liver tissues from older seals harvested from the GSL.

## ACKNOWLEDGEMENTS

We would like to thank Réjean Vigneau, Gil Thériault and members of the *Association des chasseurs de phoques Intra-Québec* (ACPIQ) for their valuable contributions to this project. Thank you to Nicolas Toupoint from MERINOV, Karine Villemaire and François Bourque from the *Ministère de l’Agriculture, des Pêcheries et de l’Alimentation du Québec* (MAPAQ), and Michael Hammill, Josée Richard and Cédric Arsenault from Fisheries and Oceans Canada (DFO) for their help with this project. Thank you to Nick Schrier (Guelph) and Dominic Bélanger (Montréal) for help with laboratory analyses. We acknowledge help with translation from French by www.DeepL.com/Translator.

## SUPPORTING INFORMATION

14 pages including data tables (S1-S15) and details on quality assurance for laboratory analyses.

