## Supplemental Information for "Trace elements in grey seals from the Gulf of St. Lawrence"

### *Supporting Information for Article: Trace elements in grey seals from the Gulf of St. Lawrence*

**CRediT AUTHOR STATEMENT. Gwyneth A. MacMillan**: Formal Analysis, Writing - Original Draft, Visualization. **Marc Amyot**: Supervision, Validation, Writing - Review & Editing. **Pierre-Yves Daoust**: Conceptualization, Methodology, Investigation, Funding Acquisition, Writing - Review & Editing. **Mélanie Lemire**: Supervision, Validation, Funding Acquisition, Writing - Review & Editing.

**Keywords :** pinnipeds, gray seal, *Halichoerus grypus*, metals, mercury, methylmercury

**Declaration of Competing Interest.** The authors declare that they have no known competing financial interests or personal relationships that influenced the work reported in this paper.

### SUPPORTING INFORMATION

**Table S1** : Methods used for trace element analysis of grey seal samples with detection limits (LOD) of the analytical instruments (in µg/g) and detection frequencies (percentage of samples > LOD).*

| | | | Detection Frequency (% > LOD) | | | | | | |
|---|---|---|---|---|---|---|---|---|---|
| | | | Group 1 | | | Group 2 | | | |
| | | | | n = 88 | n = 89 | | n = 31 | n = 31 | n = 31 |
| **Laboratory** | **Analyte** | **Method** | **LOD** | **Muscle** | **Liver** | **LOD** | **Muscle** | **Heart** | **Kidney** |
| Guelph | Antimony(Sb) | ICP-MS | 0.05 | - | - | 0.006 | - | - | 6.5 |
| | Total Arsenic (As) | ICP-MS | 0.10 | - | - | 0.004 | **100** | **100** | **100** |
| | Beryllium (Be) | ICP-MS | 0.50 | - | - | 0.003 | - | - | - |
| | Boron (B) | ICP-MS | 1.00 | - | - | 0.05 | 22.6 | 32.3 | 58.1 |
| | Cadmium (Cd) | ICP-MS | 0.05 | 1.1 | **60.7** | 0.004 | 12.9 | 22.6 | **100** |
| | Chrome (Cr) | ICP-MS | 0.50 | 27.3 | 51.7 | 0.04 | 19.4 | 16.1 | 29.0 |
| | Cobalt (Co) | ICP-MS | 0.01 | 13.6 | 55.1 | 0.006 | 3.2 | 29.0 | **100** |
| | Copper (Cu) | ICP-MS | 0.50 | **100** | **100** | 0.13 | **100** | **100** | **100** |
| | Iron(Fe) | ICP-MS | 1.00 | **100** | **100** | 2.5 | **100** | **100** | **100** |
| | Lead (Pb) | ICP-MS | 0.01 | 29.6 | 48.3 | 0.005 | 45.2 | 38.7 | **93.5** |
| | Magnesium (Mg) | ICP-MS | 2.00 | **100** | **100** | 1.2 | **100** | **100** | **100** |
| | Manganese (Mn) | ICP-MS | 0.50 | 4.5 | **100** | 0.15 | **83.9** | **100** | **100** |
| | Mercure total (Hg) | ICP-MS | 0.01 | **98.9** | **100** | 0.001 | **100** | **100** | **100** |
| | Molybdenum (Mo) | ICP-MS | 0.05 | 1.1 | **100** | 0.006 | 16.1 | **100** | **100** |
| | Nickel (Ni) | ICP-MS | 0.10 | 3.4 | - | 0.11 | - | - | 9.7 |
| | Selenium (Se) | ICP-MS | 0.05 | **100** | **100** | 0.02 | **100** | **100** | **100** |
| | Thallium (Tl) | ICP-MS | 0.005 | - | - | 0.002 | - | - | 19.4 |
| | Tin (Sn) | ICP-MS | 0.50 | - | 2.2 | 0.13 | - | - | - |
| | Zinc (Zn) | ICP-MS | 1.00 | **100** | **100** | 0.6 | **100** | **100** | **100** |
| | | | | n = 32 | n = 45 | | n = 31 | n = 31 | n = 31 |
| | | | **LOD** | **Muscle** | **Liver** | **LOD** | **Muscle** | **Heart** | **Kidney** |
| Montreal | Total Mercury (THg) | DMA-80 | 0.001 | **100** | **100** | 0.001 | **100** | **100** | **100** |
| | Total Mercury (THg) | Tekran 2600 | 0.0004 | **100** | **100** | | | | |
| | Methylmercury (MeHg) | Tekran 2700 | 0.0001 | **100** | **100** | 0.00001 | **100** | **100** | **100** |

*The symbol "-" indicates that the analyte was not detected in any of the samples. An analyte is considered "detected" if more than 60% of samples were higher than the detection limits (shown in **bold**, if detected). Sample size is shown as "n".

**Table S2** : Mean, standard deviation, coefficient of variation (CV) et percent recovery (%) for samples of certified reference material run during trace element analysis at the University of Guelph.

Group 1 (certified reference material : Tort-3, n = 13)

| Units : ppm | Ni 60 | Hg 202 | Mn 55 | Fe 57 | Co 59 | Zn 68 | As 75 | Se 78 | Mo 95 | Cd 112 | Pb 208 | V 51 | Cr 52 | Cu 63 |
|---|---|---|---|---|---|---|---|---|---|---|---|---|---|---|
| Mean | 4.96 | 0.26 | 14.43 | 159.3 | 1.01 | 125.9 | 59.22 | 10.42 | 3.06 | 40.06 | 0.19 | 8.88 | 2.21 | 454.9 |
| Standard Deviation | 0.21 | 0.01 | 0.37 | 7.29 | 0.04 | 6.95 | 2.40 | 0.31 | 0.19 | 1.05 | 0.01 | 0.45 | 0.39 | 12.11 |
| CV | 4% | 5% | 3% | 5% | 4% | 6% | 4% | 3% | 6% | 3% | 6% | 5% | 18% | 3% |
| Certified Value: | 5.3 | 0.292 | 15.6 | 179 | 1.06 | 136 | 59.5 | 10.9 | 3.44 | 42.3 | 0.225 | 9.1 | 1.95 | 497 |
| Plus/minus: | 0.24 | 0.022 | 1 | 8 | | 6 | 3.8 | 1 | 0.12 | 1.8 | 0.018 | 0.4 | 0.24 | 22 |
| % Recovery | 93% | 90% | 92% | 89% | 95% | 93% | 100% | 96% | 89% | 95% | 85% | 98% | 113% | 92% |

Group 2 (certified reference material : Tort-3, n = 3)

| Units : ppm | Ni 60 | Hg 202 | Mn 55 | Fe 57 | Co 59 | Zn 68 | As 75 | Se 78 | Mo 95 | Cd 112 | Pb 208 | V 51 | Cr 52 | Cu 63 |
|---|---|---|---|---|---|---|---|---|---|---|---|---|---|---|
| Mean | 4.95 | 0.28 | 14.38 | 165.37 | 0.95 | 127.64 | 61.08 | 9.93 | 3.24 | 39.14 | 0.19 | 8.34 | 1.77 | 456.43 |
| Standard Deviation | 0.10 | 0.02 | 0.24 | 2.19 | 0.02 | 4.18 | 0.69 | 0.25 | 0.07 | 1.16 | 0.00 | 0.16 | 0.05 | 12.75 |
| CV | 2% | 6% | 2% | 1% | 2% | 3% | 1% | 3% | 2% | 3% | 1% | 2% | 3% | 3% |
| Certified Value: | 5.3 | 0.292 | 15.6 | 179 | 1.06 | 136 | 59.5 | 10.9 | 3.44 | 42.3 | 0.225 | 9.1 | 1.95 | 497 |
| Plus/minus: | 0.24 | 0.022 | 1 | 8 | | 6 | 3.8 | 1 | 0.12 | 1.8 | 0.018 | 0.4 | 0.24 | 22 |
| % Recovery | 93% | 97% | 92% | 92% | 90% | 94% | 103% | 91% | 94% | 93% | 84% | 92% | 91% | 92% |

**Table S3** : Percent recovery (%) and standard deviation (SD) for internal controls and certified reference materials for total mercury (THg) and methylmercury (MeHg) analyses at the University of Montreal.

| **Analyte** | **Instrument** | **Tort-2** | | | **Dorm-4** | | | **Dorm-2** | | | **CALA-4** | | |
|---|---|---|---|---|---|---|---|---|---|---|---|---|---|
| Group 1 | | Recovery | SD | n | Recovery | SD | n | Recovery | SD | n | Recovery | SD | n |
| THg | DMA 80 | 99.5% | 3.8% | 4 | 92.4% | 2.6% | 3 | 102.0% | 1.6% | 4 | - | - | - |
| THg | Tekran 2600 | 98.5% | - | 1 | - | - | - | - | - | - | 106.7% | 0.8% | 4 |
| MeHg | Tekran 2700 | 96.3% | 3.4% | 9 | - | - | - | 101% | 3.3% | 5 | - | - | - |
| **Analyte** | **Instrument** | **Tort-2** | | | **Dorm-4** | | | **Dorm-2** | | | **SO-2** | | |
| Group 2 | | Recovery | SD | n | Recovery | SD | n | Recovery | SD | n | Recovery | SD | n |
| THg | DMA 80 | 102.0% | 3.4% | 19 | 104.2% | - | 1 | 96.3% | 5.5% | 10 | 103% | - | 1 |
| MeHg | Tekran 2700 | 97.9% | 3.3% | 25 | - | - | - | - | - | - | - | - | - |

**Table S4**. Inter-laboratory comparison : Total mercury concentrations (THg) measured in the same samples at two laboratories (Guelph and Montreal) by tissue and age category. Values are geometric means, minimums (min) and maximums (max). YY = Young-of-the-year.

| **Category** | **Lab** | | | | **Muscle THg** | | | | **Liver THg** |
|---|---|---|---|---|---|---|---|---|---|
| | | **n** | **Mean** | **Min** | **Max** | **n** | **Mean** | **Min** | **Max** |
| All samples (pooled) | Guelph | 119 | 0.189 | 0.005 | 1.90 | 89 | 4.67 | 0.072 | 290 |
| | Montreal | 63 | 0.312 | 0.04 | 2.26 | 45 | 5.24 | 0.140 | 454 |
| Adult | Guelph | 48 | 0.336 | 0.09 | 1.90 | 48 | 31.5 | 3.80 | 290 |
| | Montreal | 20 | 0.422 | 0.15 | 2.26 | 20 | 57.5 | 5.37 | 454 |
| Juvenile | Guelph | 17 | 0.363 | 0.15 | 0.86 | 6 | 13.80 | 8.70 | 22 |
| | Montreal | 11 | 0.459 | 0.15 | 1.22 | 6 | 17.00 | 9.91 | 30.2 |
| Spring YY | Guelph | 20 | 0.335 | 0.11 | 0.76 | | | | |
| | Montreal | 20 | 0.384 | 0.10 | 0.76 | | | | |
| Winter YY | Guelph | 34 | 0.043 | < LD | 0.18 | 35 | 0.283 | 0.072 | 2.20 |
| | Montreal | 12 | 0.093 | 0.04 | 0.17 | 19 | 0.290 | 0.14 | 0.60 |

| **Category** | **Lab** | | | | **Heart THg** | | | | **Kidney THg** |
|---|---|---|---|---|---|---|---|---|---|
| | | n | Moy. | Min | Max | n | Moy. | Min | Max |
| Spring YY and Juveniles | Guelph | 31 | 0.128 | 0.045 | 0.50 | 31 | 0.678 | 0.17 | 2.90 |
| | Montréal | 31 | 0.135 | 0.042 | 0.49 | 31 | 0.794 | 0.23 | 2.69 |

**Table S5.** Trace element concentrations detected in muscle tissues of grey seals by age category for Groups 1 and 2 (wet weight). Concentrations under the LOD were replaced with half the LOD for analytes where more than 60% of samples were detected. Values are sample size (n), geometric means, 95% confidence intervals (lower limit, upper limit), minimums (min) and maximums (max) in µg/g. YY = Young-of-the-year.

| **MUSCLE** | | | | | | | | | |
|---|---|---|---|---|---|---|---|---|---|
| **µg/g** | **LOD** | **Age Category** | **n** | **Geometric Mean** | **95% CI** | **min** | **max** | **P-value** | **Post-Hoc** |
| **Cu** | 0.50 | Total | 119 | 1.19 | (1.15, 1.23) | 0.75 | 1.8 | | |
| | 0.50 | Adult | 48 | 1.11 | (1.05, 1.18) | 0.75 | 1.8 | <0.001* | a |
| | 0.50/0.13 | Juvenile | 17 | 1.06 | (1.00, 1.12) | 0.78 | 1.2 | | a |
| | 0.13 | Spring YY | 20 | 1.33 | (1.25, 1.41) | 1.0 | 1.7 | | b |
| | 0.50 | Winter YY | 34 | 1.29 | (1.21, 1.37) | 0.82 | 1.8 | | b |
| **Fe** | 1.0 | Total | 119 | 132 | (120.56, 144.16) | 46 | 350 | | |
| | 1.0 | Adult | 48 | 166 | (152.54, 180.59) | 87 | 350 | <0.001* | a |
| | 1.0/2.5 | Juvenile | 17 | 203 | (187.18, 220.15) | 160 | 300 | | b |
| | 2.5 | Spring YY | 20 | 167 | (158.68, 175.63) | 130 | 200 | | ab |
| | 1.0 | Winter YY | 34 | 66.8 | (61.79, 72.22) | 46 | 150 | | c |
| **Pb**† | 0.010 | Total | 119 | | | < LOD | 0.48 | | |
| | 0.010 | Adult | 48 | | | < LOD | 0.48 | | |
| | 0.01/0.005 | Juvenile | 17 | | | < LOD | 0.14 | | |
| | 0.005 | Spring YY | 20 | | | < LOD | 0.097 | | |
| | 0.010 | Winter YY | 34 | | | < LOD | 0.20 | | |
| **Mg** | 2.0 | Total | 119 | 250 | (242.84, 257.48) | 160 | 380 | | |
| | 2.0 | Adult | 48 | 269 | (256.24, 282.42) | 180 | 380 | <0.001* | a |
| | 2.0/1.2 | Juvenile | 17 | 253 | (244.09, 263.01) | 220 | 300 | | a |
| | 1.2 | Spring YY | 20 | 253 | (246.13, 260.99) | 220 | 300 | | a |
| | 2.0 | Winter YY | 34 | 222 | (210.78, 234.40) | 160 | 320 | | b |
| **Se** | 0.05 | Total | 119 | 0.371 | (0.36, 0.39) | 0.20 | 0.66 | | |
| | 0.05 | Adult | 48 | 0.393 | (0.37, 0.42) | 0.24 | 0.66 | 0.0246* | a |
| | 0.05/0.02 | Juvenile | 17 | 0.378 | (0.34, 0.42) | 0.28 | 0.64 | | ab |
| | 0.02 | Spring YY | 20 | 0.371 | (0.35, 0.40) | 0.28 | 0.55 | | ab |
| | 0.05 | Winter YY | 34 | 0.340 | (0.31, 0.37) | 0.20 | 0.57 | | b |
| **Zn** | 1.00 | Total | 119 | 37.4 | (35.80, 39.10) | 18 | 92 | | |
| | 1.00 | Adult | 48 | 39.0 | (35.73, 42.66) | 18 | 92 | 0.0071* | ab |
| | 1.0/0.60 | Juvenile | 17 | 33.6 | (30.49, 37.10) | 21 | 48 | | ab |
| | 0.60 | Spring YY | 20 | 33.2 | (31.66, 34.76) | 26 | 40 | | a |
| | 1.00 | Winter YY | 34 | 39.9 | (37.71, 42.17) | 27 | 59 | | b |

* Significant P-value ($p < 0.05$), one-way ANOVA on transformed data ($log_{10}$), Post-Hoc Tukey HSD.
† Only minimum and maximum values shown because the analyte was not detected in more than 60% of samples.

**Table S6**. Trace element concentrations detected in liver tissues of grey seals by age category for Group 1 (wet weight). Concentrations under the LOD were replaced with half the LOD for analytes where more than 60% of samples were detected. Values are sample size (n), geometric means, 95% confidence intervals (lower limit, upper limit), minimums (min) and maximums (max) in µg/g. YY = Young-of-the-year.

| LIVER | | | | | | | | | |
|---|---|---|---|---|---|---|---|---|---|
| **µg/g** | **LOD** | **Age Category** | **n** | **Geometric Mean** | **95% CI** | **min** | **max** | **P-value** | **Post-Hoc** |
| **Cd** | 0.05 | Total | 54 | 1.07 | (0.88, 1.30) | < LOD | 51 | | |
| | | Adult | 48 | 1.08 | (0.87, 1.35) | 0.28 | 51 | 0.7967 | |
| | | Juvenile | 6 | 1.00 | (0.74, 1.35) | 0.57 | 1.3 | | |
| | | Winter YY | 35 | | | < LOD | < LOD | | |
| **Cr** | 0.50 | Total | 89 | 0.49 | (0.43, 0.57) | < LOD | 2.6 | | |
| | | Adult | 48 | 0.58 | (0.48, 0.72) | < LOD | 2.6 | <0.001* | a |
| | | Juvenile | 6 | 0.39 | (0.22, 0.67) | < LOD | 0.97 | | ab |
| | | Winter YY | 35 | 0.41 | (0.34, 0.50) | < LOD | 1.1 | | b |
| **Co†** | 0.01 | Total | 89 | | | < LOD | 0.027 | | |
| | | Adult | 48 | | | < LOD | 0.027 | | |
| | | Juvenile | 6 | | | < LOD | 0.025 | | |
| | | Winter YY | 35 | | | < LOD | 0.011 | | |
| **Cu** | 0.50 | Total | 89 | 28.90 | (25.93, 32.21) | 7.2 | 83 | | |
| | | Adult | 48 | 29.39 | (25.01, 34.53) | 7.2 | 72 | 0.0579 | |
| | | Juvenile | 6 | 52.75 | (41.41, 67.21) | 36 | 83 | | |
| | | Winter YY | 35 | 25.48 | (22.30, 29.10) | 13 | 57 | | |
| **Fe** | 1.00 | Total | 89 | 393.66 | (342.34, 452.68) | 65 | 1700 | | |
| | | Adult | 48 | 298.85 | (255.63, 349.37) | 65 | 1300 | <0.001* | a |
| | | Juvenile | 6 | 254.82 | (180.76, 359.21) | 150 | 530 | | a |
| | | Winter YY | 35 | 618.92 | (505.90, 757.19) | 160 | 1700 | | b |
| **Pb†** | 0.01 | Total | 89 | | | < LOD | 0.048 | | |
| | | Adult | 48 | | | < LOD | 0.048 | | |
| | | Juvenile | 6 | | | < LOD | 0.041 | | |
| | | Winter YY | 35 | | | < LOD | 0.026 | | |
| **Mg** | 2.00 | Total | 89 | 202.57 | (197.89, 207.35) | 150 | 260 | | |
| | | Adult | 48 | 190.08 | (185.26, 195.02) | 150 | 230 | <0.001* | a |
| | | Juvenile | 6 | 206.19 | (194.11, 219.02) | 180 | 220 | | ab |
| | | Winter YY | 35 | 220.36 | (214.17, 226.73) | 180 | 260 | | b |
| **Mn** | 0.50 | Total | 89 | 3.71 | (3.53, 3.90) | 2.4 | 6.4 | | |
| | | Adult | 48 | 4.07 | (3.80, 4.36) | 2.4 | 6.4 | <0.001* | a |
| | | Juvenile | 6 | 4.55 | (4.00, 5.18) | 3.6 | 5.6 | | a |
| | | Winter YY | 35 | 3.16 | (3.02, 3.31) | 2.4 | 4.6 | | b |
| **Mo** | 0.05 | Total | 89 | 0.47 | (0.45, 0.50) | 0.26 | 0.87 | | |
| | | Adult | 48 | 0.52 | (0.48, 0.56) | 0.29 | 0.85 | <0.001* | a |
| | | Juvenile | 6 | 0.62 | (0.53, 0.73) | 0.51 | 0.87 | | a |
| | | Winter YY | 35 | 0.39 | (0.37, 0.42) | 0.26 | 0.61 | | b |
| **Se** | 0.05 | Total | 89 | 4.12 | (2.97, 5.73) | 0.47 | 110 | | |
| | | Adult | 48 | 13.87 | (10.65, 18.07) | 2.2 | 110 | <0.001* | a |
| | | Juvenile | 5 | 6.28 | (4.64, 8.49) | 4.0 | 9.2 | | a |
| | | Winter YY | 34 | 0.73 | (0.67, 0.79) | 0.47 | 1.2 | | b |
| **Zn** | 1.00 | Total | 89 | 80.66 | (74.14, 87.74) | 20 | 160 | | |
| | | Adult | 48 | 65.21 | (58.51, 72.68) | 20 | 150 | <0.001* | a |
| | | Juvenile | 5 | 70.00 | (57.27, 85.57) | 55 | 110 | | a |
| | | Winter YY | 34 | 110.62 | (103.31, 118.44) | 73 | 160 | | b |

* Significant P-value ($p < 0.05$), one-way ANOVA on transformed data ($\log_{10}$), Post-Hoc Tukey HSD.
† Only minimum and maximum values shown because the analyte was not detected in more than 60% of samples.

**Table S7**. Trace element concentrations detected in muscle, heart and kidney tissues of grey seals by age category for Groups 1 (6 juveniles) and Group 2 (11 juveniles and 20 Spring YY) (wet weight). Age categories were pooled here due to the lack of significant differences between age categories for most elements. Concentrations under the LOD were replaced with half the LOD for analytes where more than 60% of samples were detected. Values are sample size (n), geometric means, 95% confidence intervals (lower limit, upper limit), minimums (min) and maximums (max) in µg/g. YY = Young-of-the-year.

| | LOD | Tissue | n | Geometric Mean | 95% CI | min | max | P-value | Post-Hoc |
|---|---|---|---|---|---|---|---|---|---|
| **As** | 0.004 | Muscle | 37 | 0.113 | (0.10, 0.13) | 0.05 | 0.28 | < 0.001* | a |
| | | Heart | 31 | 0.272 | (0.24, 0.31) | 0.17 | 0.75 | | b |
| | | Kidney | 31 | 0.238 | (0.21, 0.27) | 0.11 | 0.63 | | b |
| **Cd** | 0.004 | Muscle† | 37 | | | <LOD | 0.025 | | |
| | | Heart† | 31 | | | <LOD | 0.0068 | | |
| | | Kidney | 31 | 0.986 | (0.74, 1.31) | 0.27 | 4.3 | | |
| **Co** | 0.01 | Muscle† | 37 | | | <LOD | 0.027 | | |
| | | Heart† | 31 | | | | | | |
| | | Kidney | 31 | 0.011 | (0.01, 0.012) | <LOD | 0.026 | | |
| **Cu** | 0.50 | Muscle | 37 | 1.20 | (1.13, 1.26) | 0.78 | 1.7 | < 0.001* | a |
| | | Heart | 31 | 2.79 | (2.65, 2.93) | 1.9 | 3.4 | | b |
| | | Kidney | 31 | 2.89 | (2.77, 3.02) | 2.1 | 3.5 | | b |
| **Fe** | 1.00 | Muscle | 37 | 182.6 | (172.75, 193.07) | 130 | 300 | < 0.001* | a |
| | | Heart | 31 | 87.5 | (80.53, 94.99) | 63 | 220 | | b |
| | | Kidney | 31 | 77.3 | (71.58, 83.47) | 45 | 110 | | b |
| **Pb** | 0.01 | Muscle† | 37 | | | <LOD | 0.14 | | |
| | | Heart† | 31 | | | <LOD | 0.26 | | |
| | | Kidney | 31 | 0.017 | (0.011, 0.024) | <LOD | 0.22 | | |
| **Mg** | 2.00 | Muscle | 37 | 253.4 | (247.65, 259.32) | 220 | 300 | < 0.001* | a |
| | | Heart | 31 | 219.7 | (212.48, 227.18) | 170 | 260 | | b |
| | | Kidney | 31 | 158.3 | (153.13, 163.72) | 130 | 190 | | c |
| **Mn** | 0.50 | Muscle | 37 | 0.191 | (0.17, 0.22) | 0.075 | 0.31 | < 0.001* | a |
| | | Heart | 31 | 0.369 | (0.35, 0.39) | 0.21 | 0.5 | | b |
| | | Kidney | 31 | 0.903 | (0.85, 0.96) | 0.58 | 1.2 | | c |
| **Hg[1]** | 0.01 | Muscle | 37 | 0.348 | (0.29, 0.42) | 0.11 | 0.86 | < 0.001* | a |
| | | Heart | 31 | 0.128 | (0.10, 0.16) | 0.045 | 0.5 | | b |
| | | Kidney | 31 | 0.678 | (0.54, 0.85) | 0.17 | 2.9 | | c |
| **MeHg[2]** | 0.0001 | Muscle | 31 | 0.356 | (0.29, 0.44) | 0.10 | 1.09 | < 0.001* | a |
| | | Heart | 31 | 0.118 | (0.10, 0.15) | 0.04 | 0.45 | | b |
| | | Kidney | 31 | 0.266 | (0.21, 0.33) | 0.07 | 0.92 | | a |
| **Mo** | 0.05 | Muscle† | 37 | | | <LOD | 0.025 | < 0.001* | |
| | | Heart | 31 | 0.032 | (0.030, 0.035) | 0.023 | 0.045 | | a |
| | | Kidney | 31 | 0.117 | (0.108, 0.127) | 0.074 | 0.260 | | b |
| **Se** | 0.05 | Muscle | 37 | 0.374 | (0.35, 0.40) | 0.28 | 0.64 | < 0.001* | a |
| | | Heart | 31 | 0.439 | (0.42, 0.46) | 0.32 | 0.64 | | b |
| | | Kidney | 31 | 2.08 | (1.96, 2.21) | 1.3 | 2.7 | | c |
| **Zn** | 1.00 | Muscle | 37 | 33.4 | (31.73, 35.13) | 21 | 48 | < 0.001* | a |
| | | Heart | 31 | 26.5 | (25.55, 27.56) | 18 | 31 | | b |
| | | Kidney | 31 | 20.3 | (19.69, 20.95) | 17 | 24 | | c |

* Significant P-value ($p < 0.05$), one-way ANOVA on transformed data ($log_{10}$), Post-Hoc Tukey HSD.
†Only minimum and maximum values shown because the analyte was not detected in more than 60% of samples.
1 Analysis at the University of Guelph
2 Analysis at the University of Montréal

**Table S8**. Concentrations of total mercury (THg), methylmercury (MeHg) and the percentage of THg in the form of MeHg (% MeHg) detected in muscle and liver tissues of grey seals by age category for Groups 1 and 2 (wet weight). Concentrations under the LOD were replaced with half the LOD. Values are sample size (n), geometric means, 95% confidence intervals (lower limit, upper limit), minimums (min) and maximums (max) in µg/g. YY = Young-of-the-year.

| MUSCLE | LOD | Age Category | n | Geometric Mean | 95% CI | min | max | P-value | Post-Hoc |
|---|---|---|---|---|---|---|---|---|---|
| **THg [1]** | 0.01 | Total | 119 | 0.189 | (0.15, 0.23) | <LOD | 1.90 | | |
| | 0.01 | Adult | 48 | 0.336 | (0.28, 0.40) | 0.086 | 1.90 | <0.001* | a |
| | 0.01/0.001 | Juvenile | 17 | 0.363 | (0.27, 0.49) | 0.15 | 0.86 | | a |
| | 0.001 | Spring YY | 20 | 0.335 | (0.27, 0.42) | 0.11 | 0.76 | | a |
| | 0.01 | Winter YY | 34 | 0.043 | (0.03, 0.06) | <LOD | 0.18 | | b |
| **THg [2]** | 0.001 | Total | 63 | 0.312 | (0.25, 0.38) | 0.04 | 2.26 | | |
| | | Adult | 20 | 0.422 | (0.32, 0.56) | 0.15 | 2.26 | <0.001* | a |
| | | Juvenile | 11 | 0.459 | (0.30, 0.70) | 0.15 | 1.22 | | a |
| | | Spring YY | 20 | 0.384 | (0.31, 0.48) | 0.12 | 0.879 | | a |
| | | Winter YY | 12 | 0.093 | (0.07, 0.12) | 0.04 | 0.17 | | b |
| **MeHg [2]** | 0.0001 | Total | 63 | 0.260 | (0.21, 0.32) | 0.03 | 1.84 | | |
| | | Adult | 20 | 0.350 | (0.26, 0.47) | 0.10 | 1.84 | <0.001* | a |
| | | Juvenile | 11 | 0.400 | (0.27, 0.60) | 0.14 | 1.08 | | a |
| | | Spring YY | 20 | 0.333 | (0.27, 0.42) | 0.10 | 0.76 | | a |
| | | Winter YY | 12 | 0.071 | (0.05, 0.09) | 0.03 | 0.14 | | b |
| **% MeHg[2]** | | Total | 63 | 83.6 | (81.54, 85.77) | 62.06 | 99.85 | 0.0029* | |
| | | Adult | 20 | 83.0 | (79.97, 86.20) | 65.92 | 97.29 | | ab |
| | | Juvenile | 11 | 87.1 | (83.54, 90.90) | 77.95 | 99.85 | | a |
| | | Spring YY | 20 | 86.7 | (83.52, 90.09) | 62.06 | 95.32 | | a |
| | | Winter YY | 12 | 76.7 | (71.33, 82.48) | 63.52 | 92.96 | | b |

| LIVER | LOD | Age Category | n | Geometric Mean | 95% CI | min | max | P-value | Post-Hoc |
|---|---|---|---|---|---|---|---|---|---|
| **THg [1]** | 0.01 | Total | 89 | 4.67 | (2.79, 7.82) | 0.072 | 290 | | |
| | | Adult | 48 | 31.46 | (23.71, 41.75) | 3.8 | 290 | <0.001* | a |
| | | Juvenile | 6 | 13.80 | (9.80, 19.43) | 8.7 | 22 | | a |
| | | Winter YY | 35 | 0.283 | (0.22, 0.37) | 0.072 | 2.2 | | b |
| **THg [2]** | 0.001 | Total | 45 | 5.24 | (2.41, 11.37) | 0.14 | 454 | | |
| | | Adult | 20 | 57.51 | (35.49, 93.19) | 5.37 | 454 | <0.001* | a |
| | | Juvenile | 6 | 17.00 | (11.32, 25.54) | 9.91 | 30.2 | | b |
| | | Winter YY | 19 | 0.290 | (0.24, 0.35) | 0.14 | 0.60 | | c |
| **MeHg [2]** | 0.0001 | Total | 45 | 0.178 | (0.13, 0.24) | 0.03 | 3.22 | | |
| | | Adult | 20 | 0.349 | (0.26, 0.48) | 0.11 | 3.00 | <0.001* | a |
| | | Juvenile | 6 | 0.333 | (0.27, 0.41) | 0.25 | 0.47 | | a |
| | | Winter YY | 19 | 0.072 | (0.06, 0.09) | 0.03 | 0.17 | | b |
| **% MeHg [2]** | | Total | 45 | 3.36 | (1.93, 5.86) | 0.27 | 39.6 | | |
| | | Adult | 20 | 0.61 | (0.42, 0.87) | 0.27 | 5.16 | <0.001* | a |
| | | Juvenile | 6 | 1.97 | (1.19, 3.24) | 0.83 | 3.69 | | b |
| | | Winter YY | 19 | 24.2 | (21.1, 27.8) | 12.9 | 39.6 | | c |

* Significant P-value ($p < 0.05$), one-way ANOVA on transformed data ($\log_{10}$), Post-Hoc Tukey HSD.
1 Analysis at the University of Guelph
2 Analysis at the University of Montréal

**Table S9**. Concentrations of total mercury (THg), methylmercury (MeHg) and the percentage of THg in the form of MeHg (% MeHg) detected in heart and kidney tissues of grey seals by age category for Group 2 (wet weight). Values are sample size (n), geometric means, 95% confidence intervals (lower limit, upper limit), minimums (min) and maximums (max) in µg/g. YY = Young-of-the-year.

| HEART | LOD | Age Category | n | Geometric Mean | 95% CI | min | max | P-value |
|---|---|---|---|---|---|---|---|---|
| **THg [1]** | 0.01 | Total | 31 | 0.128 | (0.10, 0.16) | 0.045 | 0.50 | |
| | | Juvenile | 11 | 0.139 | (0.09, 0.22) | 0.045 | 0.50 | 0.5639 |
| | | Spring YY | 20 | 0.122 | (0.10, 0.15) | 0.045 | 0.27 | |
| **THg [2]** | 0.001 | Total | 31 | 0.135 | (0.11, 0.17) | 0.042 | 0.486 | |
| | | Juvenile | 11 | 0.148 | (0.10, 0.23) | 0.052 | 0.486 | 0.5379 |
| | | Spring YY | 20 | 0.129 | (0.10, 0.16) | 0.042 | 0.32 | |
| **MeHg [2]** | 0.0001 | Total | 31 | 0.118 | (0.10, 0.15) | 0.037 | 0.448 | |
| | | Juvenile | 11 | 0.129 | (0.08, 0.20) | 0.050 | 0.45 | 0.5946 |
| | | Spring YY | 20 | 0.113 | (0.09, 0.14) | 0.037 | 0.30 | |
| **% MeHg[2]** | | Total | 31 | 87.7 | (86.05, 89.36) | 76.94 | 96.61 | |
| | | Juvenile | 11 | 86.9 | (83.56, 90.34) | 76.94 | 94.98 | 0.4862 |
| | | Spring YY | 20 | 88.1 | (86.35, 89.95) | 81.71 | 96.61 | |

| KIDNEY | LOD | Age Category | n | Geometric Mean | 95% CI | min | max | P-value |
|---|---|---|---|---|---|---|---|---|
| **THg [1]** | 0.01 | Total | 31 | 0.678 | (0.54, 0.85) | 0.17 | 2.90 | |
| | | Juvenile | 11 | 0.853 | (0.54, 1.34) | 0.39 | 2.90 | 0.1422 |
| | | Spring YY | 20 | 0.597 | (0.47, 0.76) | 0.17 | 1.30 | |
| **THg [2]** | 0.001 | Total | 31 | 0.794 | (0.64, 0.99) | 0.228 | 2.69 | |
| | | Juvenile | 11 | 1.08 | (0.72, 1.64) | 0.499 | 2.69 | 0.0357* ꝉ |
| | | Spring YY | 20 | 0.669 | (0.54, 0.84) | 0.228 | 1.59 | |
| **MeHg [2]** | 0.0001 | Total | 31 | 0.266 | (0.21, 0.33) | 0.072 | 0.918 | |
| | | Juvenile | 11 | 0.285 | (0.18, 0.45) | 0.082 | 0.918 | 0.650 |
| | | Spring YY | 20 | 0.255 | (0.20, 0.32) | 0.072 | 0.612 | |
| **% MeHg[2]** | | Total | 31 | 33.4 | (30.41, 36.76) | 16.4 | 53.0 | |
| | | Juvenile | 11 | 26.3 | (22.48, 30.79) | 16.4 | 38.2 | < 0.001* |
| | | Spring YY | 20 | 38.2 | (35.58, 40.91) | 30.1 | 53.0 | |

* Significant P-value ($p < 0.05$), one-way ANOVA on transformed data ($\log_{10}$), Post-Hoc Tukey HSD.

1 Analysis at the University of Guelph

2 Analysis at the University of Montreal

ꝉ Note the significant difference between juveniles and Spring YY for kidney samples analysed at the University of Montreal, but no difference for samples analysed at the University of Guelph. This may be due to the small sample size (n = 11) and the presence of an outlier in the sub-set of samples analysed in Montreal.

**Table S10.** Trace element concentrations detected in muscle tissues of grey seals by sex for Group 1 (adults and juveniles only) (wet weight). Concentrations under the LOD were replaced with half the LOD for analytes where more than 60% of samples were detected. Values are sample size (n), geometric means, 95% confidence intervals (lower limit, upper limit), minimums (min) and maximums (max) in µg/g.

| MUSCLE | | | | | | | | |
|---|---|---|---|---|---|---|---|---|
| **µg/g** | **LOD** | **Sex** | **n** | **Geometric Mean** | **95% CI** | **min** | **max** | **P-value** |
| **Cu** | 0.50 | Total | 65 | 1.10 | (1.05, 1.15) | 0.75 | 1.8 | |
| | | Female | 19 | 1.06 | (0.99, 1.14) | 0.82 | 1.5 | 0.3390 |
| | | Male | 46 | 1.11 | (1.05, 1.18) | 0.75 | 1.8 | |
| **Fe** | 1.00 | Total | 65 | 175 | (163.28, 187.45) | 87 | 350 | |
| | | Female | 19 | 176 | (157.58, 196.50) | 110 | 300 | 0.9165 |
| | | Male | 46 | 175 | (159.99, 190.38) | 87 | 350 | |
| **Pb** | 0.01 | Total | 65 | | | < LOD | 0.48 | |
| | | Female | 19 | | | < LOD | 0.26 | |
| | | Male | 46 | | | < LOD | 0.48 | |
| **Mg** | 2.00 | Total | 65 | 265 | (255.05, 274.99) | 180 | 380 | |
| | | Female | 19 | 259 | (244.82, 273.32) | 210 | 320 | 0.4359 |
| | | Male | 46 | 267 | (254.84, 280.60) | 180 | 380 | |
| **Hg[1]** | 0.01 | Total | 65 | 0.343 | (0.29, 0.40) | 0.086 | 1.9 | |
| | | Female | 19 | 0.246 | (0.20, 0.30) | 0.14 | 0.59 | 0.0062* |
| | | Male | 46 | 0.394 | (0.32, 0.48) | 0.086 | 1.9 | |
| **MeHg[2]** | 0.0001 | Total | 65 | 0.367 | (0.29, 0.46) | 0.10 | 1.84 | |
| | | Female | 19 | 0.247 | (0.19, 0.33) | 0.13 | 0.66 | 0.0323* |
| | | Male | 46 | 0.431 | (0.32, 0.57) | 0.10 | 1.84 | |
| **Se** | 0.05 | Total | 65 | 0.389 | (0.37, 0.41) | 0.24 | 0.66 | |
| | | Female | 19 | 0.373 | (0.35, 0.40) | 0.29 | 0.45 | 0.2781 |
| | | Male | 46 | 0.396 | (0.37, 0.42) | 0.24 | 0.66 | |
| **Zn** | 1.00 | Total | 65 | 37.5 | (34.95, 40.34) | 18 | 92 | |
| | | Female | 19 | 36.7 | (32.25, 41.67) | 18 | 65 | 0.6779 |
| | | Male | 46 | 37.9 | (34.76, 41.37) | 21 | 92 | |

* Significant P-value ($p < 0.05$), independent sample Student's t-test, on transformed data ($log_{10}$).
1 Analysis at the University of Guelph
2 Analysis at the University of Montréal

**Table S11.** Trace element concentrations detected in liver tissues of grey seals by sex for Group 1 (adults and juveniles only) (wet weight). Concentrations under the LOD were replaced with half the LOD for analytes where more than 60% of samples were detected. Values are sample size (n), geometric means, 95% confidence intervals (lower limit, upper limit), minimums (min) and maximums (max) in µg/g.

| LIVER | LOD | Sex | n | Geometric Mean | 95% CI | min | max | P-value |
|---|---|---|---|---|---|---|---|---|
| **Cd** | 0.05 | Total | 54 | 1.07 | (0.89, 1.30) | 0.28 | 51 | |
| | | Female | 14 | 1.27 | (0.71, 2.27) | 0.57 | 51 | 0.3138 |
| | | Male | 40 | 1.01 | (0.86, 1.19) | 0.28 | 2.3 | |
| **Cr** | 0.50 | Total | 54 | 0.559 | (0.46, 0.68) | 0.25 | 2.6 | |
| | | Female | 14 | 0.501 | (0.34, 0.73) | 0.25 | 1.3 | 0.5153 |
| | | Male | 40 | 0.581 | (0.46, 0.73) | 0.25 | 2.6 | |
| **Co** | 0.01 | Total | 54 | 0.013 | (0.0114, 0.0142) | < LOD | 0.027 | |
| | | Female | 14 | 0.014 | (0.0114, 0.0178) | < LOD | 0.027 | 0.2443 |
| | | Male | 40 | 0.012 | (0.0108, 0.0139) | < LOD | 0.025 | |
| **Cu** | 0.50 | Total | 54 | 31.4 | (26.89, 36.57) | 7.2 | 83 | |
| | | Female | 14 | 40.0 | (31.59, 50.64) | 11 | 65 | 0.0659 |
| | | Male | 40 | 28.8 | (23.95, 34.64) | 7.2 | 83 | |
| **Fe** | 1.00 | Total | 54 | 294 | (254.30, 338.97) | 65 | 1300 | |
| | | Female | 14 | 244 | (178.19, 333.00) | 65 | 730 | 0.1332 |
| | | Male | 40 | 313 | (267.71, 366.95) | 100 | 1300 | |
| **Pb** | 0.01 | Total | 54 | 0.0101 | (0.0083, 0.0122) | 0.005 | 0.048 | |
| | | Female | 14 | 0.0124 | (0.0081, 0.0189) | 0.005 | 0.048 | 0.2168 |
| | | Male | 40 | 0.0094 | (0.0076, 0.0116) | 0.005 | 0.034 | |
| **Mg** | 2.00 | Total | 54 | 192 | (187.15, 196.58) | 150 | 230 | |
| | | Female | 14 | 193 | (182.52, 203.34) | 150 | 220 | 0.8387 |
| | | Male | 40 | 192 | (186.28, 196.90) | 150 | 230 | |
| **Mn** | 0.50 | Total | 54 | 4.12 | (3.87, 4.39) | 2.4 | 6.4 | |
| | | Female | 14 | 3.44 | (2.99, 3.95) | 2.4 | 5.6 | <0.001* |
| | | Male | 40 | 4.39 | (4.14, 4.66) | 2.9 | 6.4 | |
| **Hg**[1] | 0.01 | Total | 54 | 28.7 | (22.07, 37.35) | 3.8 | 290 | |
| | | Female | 14 | 35.0 | (22.05, 55.48) | 8.7 | 140 | 0.3890 |
| | | Male | 40 | 26.8 | (19.52, 36.77) | 3.8 | 290 | |
| **MeHg**[2] | 0.0001 | Total | 54 | 0.345 | (0.27, 0.44) | 0.11 | 3.22 | |
| | | Female | 14 | 0.300 | (0.25, 0.36) | 0.21 | 0.41 | 0.5045 |
| | | Male | 40 | 0.364 | (0.26, 0.50) | 0.11 | 3.22 | |
| **Mo** | 0.05 | Total | 54 | 0.530 | (0.50, 0.57) | 0.29 | 0.87 | |
| | | Female | 14 | 0.519 | (0.45, 0.60) | 0.32 | 0.8 | 0.7341 |
| | | Male | 40 | 0.533 | (0.49, 0.58) | 0.29 | 0.87 | |
| **Se** | 0.05 | Total | 54 | 12.7 | (9.93, 16.25) | 2.2 | 110 | |
| | | Female | 14 | 15.4 | (9.84, 24.05) | 4 | 64 | 0.3713 |
| | | Male | 40 | 11.9 | (8.85, 15.93) | 2.2 | 110 | |
| **Zn** | 1.00 | Total | 54 | 65.7 | (59.55, 72.54) | 20 | 150 | |
| | | Female | 14 | 51.7 | (43.22, 61.89) | 20 | 76 | 0.0039* |
| | | Male | 40 | 71.5 | (64.23, 79.54) | 32 | 150 | |

* Significant P-value ($p < 0.05$), independent sample Student's t-test, on transformed data ($\log_{10}$).
1 Analysis at the University of Guelph
2 Analysis at the University of Montréal

**Table S12**. Trace element concentrations detected in muscle tissues of grey seals by sample site for Group 1 (adults and juveniles only) (wet weight). Concentrations under the LOD were replaced with half the LOD for analytes where more than 60% of samples were detected. Values are sample size (n), geometric means, 95% confidence intervals (lower limit, upper limit), minimums (min) and maximums (max) in µg/g.

| MUSCLE | | | | | | | | | |
|---|---|---|---|---|---|---|---|---|---|
| **µg/g** | **LOD** | **Site** | **n** | **Geometric Mean** | **95% CI** | **min** | **max** | **P-value** | **Post-Hoc** |
| **Cu** | 0.50 | 1 | 27 | 1.07 | 1.07 (1.01, 1.14) | 0.78 | 1.5 | 0.0057* | a |
| | | 2 | 15 | 1.06 | 1.06 (1.01, 1.12) | 0.87 | 1.2 | | a |
| | | 3 | 8 | 0.98 | 0.98 (0.88, 1.10) | 0.81 | 1.3 | | a |
| | | 4 | 15 | 1.26 | 1.26 (1.11, 1.42) | 0.75 | 1.8 | | b |
| **Fe** | 1.00 | 1 | 27 | 161 | 161.34 (146.00, 178.28) | 87 | 240 | 0.1144 | |
| | | 2 | 15 | 181 | 181.12 (155.19, 211.40) | 93 | 300 | | |
| | | 3 | 8 | 168 | 167.82 (154.92, 181.79) | 130 | 190 | | |
| | | 4 | 15 | 200 | 199.88 (169.72, 235.41) | 120 | 350 | | |
| **Pb†** | 0.01 | 1 | 27 | | | < LOD | 0.48 | | |
| | | 2 | 15 | | | < LOD | 0.30 | | |
| | | 3 | 8 | | | < LOD | < LOD | | |
| | | 4 | 15 | | | < LOD | 0.15 | | |
| **Mg** | 2.00 | 1 | 27 | 276 | (263.54, 289.81) | 220 | 370 | 0.0028* | a |
| | | 2 | 15 | 252 | (238.43, 266.26) | 200 | 300 | | ab |
| | | 3 | 8 | 228 | (210.84, 245.60) | 190 | 260 | | b |
| | | 4 | 15 | 280 | (252.71, 309.18) | 180 | 380 | | a |
| **Hg** | 0.01 | 1 | 27 | 0.324 | (0.26, 0.41) | 0.086 | 0.86 | 0.0073* | a |
| | | 2 | 15 | 0.257 | (0.20, 0.34) | 0.13 | 0.64 | | a |
| | | 3 | 8 | 0.299 | (0.20, 0.46) | 0.14 | 0.94 | | ab |
| | | 4 | 15 | 0.547 | (0.40, 0.74) | 0.25 | 1.9 | | b |
| **MeHg** | 0.0001 | 1 | 27 | 0.400 | (0.27, 0.60) | 0.135 | 1.085 | 0.3583 | |
| | | 2 | 15 | na | na | na | na | | |
| | | 3 | 8 | 0.273 | (0.17, 0.45) | 0.1 | 1.06 | | |
| | | 4 | 15 | 0.412 | (0.29, 0.58) | 0.19 | 1.84 | | |
| **Se** | 0.05 | 1 | 27 | 0.357 | (0.34, 0.38) | 0.28 | 0.45 | 0.0160* | b |
| | | 2 | 15 | 0.403 | (0.37, 0.44) | 0.32 | 0.64 | | ab |
| | | 3 | 8 | 0.395 | (0.37, 0.42) | 0.34 | 0.44 | | ab |
| | | 4 | 15 | 0.435 | (0.38, 0.50) | 0.24 | 0.66 | | a |
| **Zn** | 1.00 | 1 | 27 | 33.9 | (31.43, 36.53) | 24 | 48 | <0.001* | a |
| | | 2 | 15 | 32.7 | (28.47, 37.61) | 18 | 43 | | a |
| | | 3 | 8 | 50.3 | (44.01, 57.52) | 36 | 65 | | b |
| | | 4 | 15 | 44.3 | (37.48, 52.45) | 27 | 92 | | b |

* Significant P-value ($p < 0.05$), one-way ANOVA on transformed data ($\log_{10}$), Post-Hoc Tukey HSD.
† Only minimum and maximum values shown because the analyte was not detected in more than 60% of samples.

**Table S13.** Trace element concentrations detected in liver tissues of grey seals by sample site for Group 1 (adults and juveniles only) (wet weight). Concentrations under the LOD were replaced with half the LOD for analytes where more than 60% of samples were detected. Values are sample size (n), geometric means, 95% confidence intervals (lower limit, upper limit), minimums (min) and maximums (max) in µg/g.

| LIVER | LD | Site | n | Geometric Mean | 95% CI | min | max | P-value | Post-Hoc |
|---|---|---|---|---|---|---|---|---|---|
| **Cd** | 0.05 | 1 | 16 | 1.02 | (0.86, 1.20) | 0.57 | 2.0 | 0.3028 | |
| | | 2 | 15 | 1.02 | (0.83, 1.26) | 0.54 | 2.3 | | |
| | | 3 | 8 | 1.67 | (0.63, 4.41) | 0.81 | 51 | | |
| | | 4 | 15 | 0.942 | (0.65, 1.37) | 0.28 | 2.2 | | |
| **Cr** | 0.50 | 1 | 16 | 0.944 | (0.92, 0.97) | 0.86 | 1.0 | <0.001* | b |
| | | 2 | 15 | 0.250 | (0.25, 0.25) | 0.25 | 0.25 | | a |
| | | 3 | 8 | 0.250 | (0.25, 0.25) | 0.25 | 0.25 | | a |
| | | 4 | 15 | 1.10 | (0.95, 1.27) | 0.83 | 2.6 | | c |
| **Co†** | 0.01 | 1 | 16 | | | 0.01 | 0.022 | | |
| | | 2 | 15 | | | < LOD | 0.025 | | |
| | | 3 | 8 | | | < LOD | 0.022 | | |
| | | 4 | 15 | | | < LOD | 0.027 | | |
| **Cu** | 0.50 | 1 | 16 | 30.0 | (22.63, 39.71) | 9.5 | 72 | 0.0328* | ab |
| | | 2 | 15 | 44.6 | (37.79, 52.58) | 28 | 83 | | a |
| | | 3 | 8 | 26.4 | (15.49, 45.06) | 8.7 | 61 | | ab |
| | | 4 | 15 | 25.4 | (19.22, 33.47) | 7.2 | 65 | | b |
| **Fe** | 1.00 | 1 | 16 | 240 | (200.48, 287.09) | 100 | 530 | <0.001* | a |
| | | 2 | 15 | 230 | (173.81, 303.67) | 65 | 720 | | a |
| | | 3 | 8 | 254 | (188.44, 341.76) | 150 | 440 | | a |
| | | 4 | 15 | 503 | (420.66, 601.57) | 350 | 1300 | | b |
| **Pb†** | 0.01 | 1 | 16 | | | < LOD | 0.048 | | |
| | | 2 | 15 | | | < LOD | 0.022 | | |
| | | 3 | 8 | | | < LOD | 0.017 | | |
| | | 4 | 15 | | | < LOD | 0.034 | | |
| **Mg** | 2.00 | 1 | 16 | 189 | (181.23, 197.75) | 150 | 220 | 0.115 | |
| | | 2 | 15 | 199 | (191.84, 206.14) | 180 | 230 | | |
| | | 3 | 8 | 197 | (192.57, 202.33) | 190 | 210 | | |
| | | 4 | 15 | 185 | (173.85, 196.34) | 150 | 220 | | |
| **Mn** | 0.50 | 1 | 16 | 3.97 | (3.49, 4.51) | 2.4 | 5.7 | 0.3558 | |
| | | 2 | 15 | 4.48 | (4.10, 4.89) | 2.8 | 5.5 | | |
| | | 3 | 8 | 3.81 | (3.25, 4.46) | 3.2 | 5.9 | | |
| | | 4 | 15 | 4.12 | (3.61, 4.69) | 2.6 | 6.4 | | |
| **Hg** | 0.01 | 1 | 16 | 23.9 | (14.42, 39.59) | 3.8 | 140 | 0.0034* | a |
| | | 2 | 15 | 16.3 | (12.40, 21.48) | 6.0 | 46 | | a |
| | | 3 | 8 | 34.3 | (15.58, 75.62) | 6.2 | 150 | | ab |
| | | 4 | 15 | 55.9 | (36.30, 85.94) | 6.4 | 290 | | b |
| **MeHg** | 0.0001 | 1 | 16 | 0.373 | (0.31, 0.45) | 0.34 | 0.41 | 0.2564 | |
| | | 2 | 15 | 0.314 | (0.24, 0.42) | 0.25 | 0.47 | | |
| | | 3 | 8 | 0.247 | (0.16, 0.38) | 0.11 | 0.79 | | |
| | | 4 | 15 | 0.440 | (0.30, 0.65) | 0.24 | 3.22 | | |
| **Mo** | 0.05 | 1 | 16 | 0.501 | (0.45, 0.55) | 0.32 | 0.67 | 0.3317 | |
| | | 2 | 15 | 0.583 | (0.52, 0.66) | 0.36 | 0.87 | | |
| | | 3 | 8 | 0.501 | (0.43, 0.58) | 0.36 | 0.72 | | |
| | | 4 | 15 | 0.526 | (0.45, 0.61) | 0.29 | 0.79 | | |
| **Se** | 0.05 | 1 | 16 | 11.0 | (6.86, 17.64) | 2.2 | 64 | 0.0049* | ab |
| | | 2 | 15 | 7.53 | (5.95, 9.52) | 3.4 | 21 | | b |
| | | 3 | 8 | 14.3 | (6.71, 30.56) | 2.8 | 60 | | ab |
| | | 4 | 15 | 23.4 | (15.39, 35.69) | 2.7 | 110 | | a |
| **Zn** | 1.00 | 1 | 16 | 54.3 | (46.31, 63.57) | 20 | 75 | <0.001* | ab |
| | | 2 | 15 | 67.5 | (61.16, 74.54) | 50 | 110 | | a |
| | | 3 | 8 | 45.1 | (37.52, 54.31) | 32 | 75 | | b |
| | | 4 | 15 | 95.9 | (85.10, 108.12) | 71 | 150 | | c |

* Significant P-value ($p < 0.05$), independent sample Student's t-test, on transformed data ($\log_{10}$).
†Only minimum and maximum values shown because the analyte was not detected in more than 60% of samples.

**Table S14 :** Correlations of trace element concentrations ($log_{10}$ ug/g) and % MeHg versus the exact age of grey seals, determined by the layers of dental cementum in the canines. The values in the table are Pearson's r correlation coefficients and significant relationships are show in bold ($p > 0.05$). Sample size (n) is shown and the asterisk shows exceptions. n.d.: not detected.

| Tissue | Age | n | Cd | Cr | Cu | Fe | Mg | Mn | Hg | MeHg* | %MeHg* | Mo | Se | Zn |
|---|---|---|---|---|---|---|---|---|---|---|---|---|---|---|
| Muscle | All categories | 54 | n.d. | n.d. | -0.12 | 0.18 | 0.20 | n.d. | 0.19 | -0.05 | -0.32 | n.d. | 0.33 | **0.60** |
| Muscle | Adult seals | 23 | n.d. | n.d. | 0.17 | 0.02 | 0.27 | n.d. | 0.03 | -0.21 | -0.30 | n.d. | -0.06 | 0.10 |
| Liver | Adult seals | 23 | -0.03 | 0.17 | -0.03 | -0.17 | -0.15 | -0.29 | -0.30 | -0.24 | 0.36 | -0.27 | -0.30 | 0.03 |

* Sample sizes for MeHg et % MeHg are n =51, n = 20 and n = 20 respectively for the 3 columns.

**Table S15**. Weekly Intake Scenarios: Percent Dietary Reference Intakes (% DRI) for grey seal muscle and liver tissues from adult seals (> 4 years) or winter young-of-the-year (winter YY, < 6 weeks). Four consumption scenarios were created based on Recommended Dietary Allowances (RDA) and Tolerable Upper Limits (UL) for two vulnerable population subgroups, 75 g of seal muscle or liver per week for children 1 – 3 years old and 150 g of seal muscle or liver per week for pregnant women 19 – 30 years old. %DRI higher than 70% are shown in **bold**.

| **Recommendations** | **DRI** | | **Population Group** | | | | **Units** | **Cr** | **Cu** | **Fe** | **Mg** | **Mn** | **Mo** | **Se** | **Zn** |
|---|---|---|---|---|---|---|---|---|---|---|---|---|---|---|---|
| | RDA | | Children 1-3 y | | | | µg/day | 11* | 340 | 7000 | 80000 | 1200* | 17 | 20 | 3000 |
| | | | Pregnant 19-30 y | | | | µg/day | 30* | 1000 | 27000 | 350000 | 2000* | 50 | 60 | 11000 |
| | UL | | Children 1-3 y | | | | µg/day | - | 1000 | 40000 | 65000† | 2000 | 300 | 90 | 7000 |
| | | | Pregnant 19-30 y | | | | µg/day | - | 10000 | 45000 | 350000† | 11000 | 2000 | 400 | 40000 |
| **Concentrations** | | | | **Seal Age** | **Tissue** | | **Units** | **Cr** | **Cu** | **Fe** | **Mg** | **Mn** | **Mo** | **Se** | **Zn** |
| | | | | Adult | Muscle | | µg /g | - | 1.11 | 166 | 269 | - | - | 0.393 | 39.0 |
| | | | | Winter YY | Muscle | | µg /g | - | 1.29 | 66.8 | 222 | - | - | 0.34 | 39.9 |
| | | | | Adult | Liver | | µg /g | 0.585 | 29.4 | 299 | 190 | 4.07 | 0.520 | 13.87 | 65.2 |
| | | | | Winter YY | Liver | | µg /g | 0.410 | 25.5 | 619 | 220 | 3.16 | 0.393 | 0.727 | 111 |
| **Estimated Intake** | **% DRI** | **Scenario** | **Population Group** | **Seal Age** | **Tissue** | **Portion** | **Units** | **Cr** | **Cu** | **Fe** | **Mg** | **Mn** | **Mo** | **Se** | **Zn** |
| | % RDA | 1) | Children 1-3 y | Adult | Muscle | 75g/week | % | - | 3.5 | 25 | 3.6 | - | - | 21 | 14 |
| | | 2) | | Winter YY | Muscle | 75g/week | % | - | 4.1 | 10 | 3.0 | - | - | 18 | 14 |
| | | 3) | | Adult | Liver | 75g/week | % | 57 | **93** | 46 | 2.5 | 3.6 | 33 | **742** | 23 |
| | | 4) | | Winter YY | Liver | 75g/week | % | 40 | **80** | **95** | 2.9 | 2.8 | 25 | 39 | 40 |
| | | 1) | Pregnant 19-30 y | Adult | Muscle | 150g/week | % | - | 2.4 | 13 | 1.6 | - | - | 14 | 7.6 |
| | | 2) | | Winter YY | Muscle | 150g/week | % | - | 2.8 | 5.3 | 1.4 | - | - | 12 | 7.8 |
| | | 3) | | Adult | Liver | 150g/week | % | 42 | 63 | 24 | 1.2 | 4.4 | 22 | **495** | 13 |
| | | 4) | | Winter YY | Liver | 150g/week | % | 29 | 55 | 49 | 1.3 | 3.4 | 17 | 26 | 22 |
| | % UL | 1) | Children 1-3 y | Adult | Muscle | 75g/week | % | - | 1.2 | 4.4 | 4.4 | - | - | 4.7 | 6.0 |
| | | 2) | | Winter YY | Muscle | 75g/week | % | - | 1.4 | 1.8 | 3.7 | - | - | 4.0 | 6.1 |
| | | 3) | | Adult | Liver | 75g/week | % | - | 32 | 8.0 | 3.1 | 2.2 | 1.9 | **165** | 10 |
| | | 4) | | Winter YY | Liver | 75g/week | % | - | 27 | 17 | 3.6 | 1.7 | 1.4 | 8.7 | 17 |
| | | 1) | Pregnant 19-30 y | Adult | Muscle | 150g/week | % | - | 0.2 | 7.9 | 1.6 | - | - | 2.1 | 2.1 |
| | | 2) | | Winter YY | Muscle | 150g/week | % | - | 0.3 | 3.2 | 1.4 | - | - | 1.8 | 2.1 |
| | | 3) | | Adult | Liver | 150g/week | % | - | 6.3 | 14 | 1.2 | 0.8 | 0.6 | **74** | 3.5 |
| | | 4) | | Winter YY | Liver | 150g/week | % | - | 5.5 | 29 | 1.3 | 0.6 | 0.4 | 3.9 | 5.9 |

**Source:** Food and Nutrition Board, Institute of Medicine, National Academy of Sciences, 2011: https://ods.od.nih.gov/Health_Information/Dietary_Reference_Intakes.aspx.

**Notes**: Recommended Dietary Allowances (RDAs) are shown in ordinary type and Adequate Intakes (AIs) are followed by an asterisk (*). An RDA is the average daily dietary intake level sufficient to meet the nutrient requirements of nearly all (97–98 percent) healthy individuals in a group. If sufficient scientific evidence is not available to calculate an RDA, an AI is usually developed. The AI is believed to cover the needs of all healthy individuals in the groups, but lack of data or uncertainty in the data prevent being able to specify with confidence the percentage of individuals covered by this intake. A Tolerable Upper Intake Level (UL) is the highest level of daily nutrient intake that is likely to pose no risk of adverse health effects to almost all individuals in the general population. Unless otherwise specified, the UL represents total intake from food, water, and supplements. Members of the general population should be advised not to routinely exceed the UL. The UL is not meant to apply to individuals who are treated with the nutrient under medical supervision or to individuals with predisposing conditions that modify their sensitivity to the nutrient.
* Adequate Intakes (AIs), where sufficient scientific evidence is not available to establish and RDA.
† The ULs for magnesium represent intake from a pharmacological agent only and do not include intake from food and water.
